# WEE1 inhibitors synergise with mRNA translation defects via activation of the kinase GCN2

**DOI:** 10.1101/2025.03.12.642825

**Authors:** Jordan C. J. Wilson, JiaYi Zhu, Simon Lam, Alexandra Hart, Chen Gang Goh, Fadia Bou-Dagher, Hlib Razumkov, Lena Kobel, Zacharias Kontarakis, John Fielden, Moritz F. Schlapansky, Joanna I. Loizou, Andreas Villunger, Jacob E. Corn, Stefan J. Marciniak, Aldo S. Bader, Stephen P. Jackson

**Affiliations:** Cancer Research UK Cambridge Institute, University of Cambridge, Cambridge, United Kingdom; Center for Cancer Research, Comprehensive Cancer Centre, Medical University of Vienna, Austria; CeMM Research Center for Molecular Medicine of the Austrian Academy of Sciences, Vienna, Austria; Cambridge Institute for Medical Research (CIMR), University of Cambridge, Cambridge, United Kingdom; Department of Chemistry, Stanford School of Humanities and Sciences, Stanford University, Stanford, California 94305, United States; Department of Chemical and Systems Biology, ChEM-H, Stanford School of Medicine, Stanford University, Stanford, California 94305, United States; Department of Biology, Institute of Molecular Health Sciences, ETH Zurich, Zurich, Switzerland; Genome Engineering and Measurement Lab, ETH Zurich, Zurich, Switzerland; Breast Cancer Now Toby Robins Research Centre, The Institute of Cancer Research, London, United Kingdom; Institute of Developmental Immunology, Biocenter, Medical University of Innsbruck, Innsbruck, Austria

## Abstract

Inhibitors of the protein kinase WEE1 have emerged as promising agents for cancer therapy. In this study, we uncover synergistic interactions between WEE1 small-molecule inhibitors and defects in mRNA translation, mediated by activation of the integrated stress response (ISR) through the kinase GCN2. Using a pooled CRISPRi screen, we identify GSPT1 and ALKBH8 as factors whose depletion confer hypersensitivity to the WEE1 inhibitor, AZD1775. We demonstrate that this synergy depends on ISR activation, which is induced by the off-target activity of WEE1 inhibitors. Furthermore, PROTAC-based WEE1 inhibitors and molecular glues show reduced or no ISR activation, suggesting potential strategies to minimise off-target toxicity. Our findings reveal that certain WEE1 inhibitors elicit dual toxicity via ISR activation and genotoxic stress, with ISR activation being independent of WEE1 itself or cell-cycle status. This dual mechanism highlights opportunities for combination therapies, such as pairing WEE1 inhibitors with agents targeting the mRNA translation machinery. This study also underscores the need for more precise WEE1 targeting strategies to mitigate off-target effects, with implications for optimising the therapeutic potential of WEE1 inhibitors.

## INTRODUCTION

WEE1 inhibition is attracting substantial interest as a target in cancer therapy, with several clinical trials using AZD1775, Zn-c3 or Debio0123 as a small molecule inhibitor of WEE1 (ClinicalTrials.gov: NCT03668340, NCT04439227, NCT05128825, NCT05743036 and NCT03968653). Furthermore, recently developed WEE1 inhibitors that include ACR-2316, IMP7068 and SY-4835 are due to undergo trials to test their safety and efficacy (ClinicalTrials.gov: NCT06667141, NCT04768868 and NCT05291182). The rationale behind the use of WEE1 inhibitors is to both increase genotoxicity in S phase of the cell cycle and to override the cell cycle G2-M checkpoint through CDK1 and CDK2 overactivation.

WEE1 serves as an essential negative regulator of cyclin-dependent kinases CDK1 and CDK2 through tyrosine phosphorylation^1,2,3,4,5^. Inhibition of WEE1 resulting in overactivation of CDK1 and CDK2 can impact on several key cellular processes. These include impacting cell-cycle checkpoint control^6^, replication origin activity^7^, control over nucleotide pools via stabilisation of the ribonucleotide reductase subunit RRM2^8^ and the protection of stalled replication forks^9^.

To enhance the efficacy of WEE1 inhibitors and mitigate toxicity, previous studies have explored predictive biomarkers and combination strategies. WEE1 inhibitors have been noted to exhibit increased sensitivity in various genetic backgrounds including loss of function TP53 with oncogenic KRAS mutations^10^, deficiencies in SIRT1^11^, ATRX^12^, RBM10^13^, FBH1^14^, as well as a reduction of the histone mark H3K36me3^8^. Despite attempts for patient stratification in the use of WEE1 inhibitors, their utility in the clinic has been limited due to several associated toxicities, including neutropenia, thrombocytopenia, nausea and anaemia^15,16^.

In this study, we uncover factors involved in mRNA translation that confer hypersensitivity to the WEE1 inhibitors. We find that this synergy is predicated on the ability of WEE1 small-molecule inhibitors to activate the integrated stress response (ISR) via the activity of the translation initiation factor eIF2α kinase, GCN2. The ISR is an evolutionarily conserved cellular signalling pathway that is initiated by the phosphorylation of the translation initiation factor eIF2α on Ser-51^17^, leading to attenuation of bulk protein synthesis and reprogramming of gene expression. This process is mediated by eIF2α kinases, including GCN2^18^. Our experiments show that several WEE1 small-molecule inhibitors activate the integrated stress response in various cancer and non-cancer cell lines via off-target ISR toxicity. Additionally, we note that WEE1 PROTAC degraders, which utilise WEE1 small-molecule inhibitors as warheads, elicit less ISR toxicity, while WEE1 molecular glues do not activate this pathway.

## RESULTS

### CRISPR screen connects WEE1 inhibitors with mRNA translation defects

To identify novel genes associated with sensitivity or resistance to WEE1 inhibitors, we performed a pooled CRISPR interference (CRISPRi) screen in the untransformed, immortalised cell line RPE-1 inactivated for *TP53* and expressing the transcriptional repressor dCas9-KRAB. This work used a DNA damage response (DDR) and cell cycle focused library targeting over 2000 genes for transcriptional repression to identify CRISPR single guide RNAs (sgRNAs) that either increased or decreased relative cell viability in response to treatment with the WEE1 inhibitor, Adavosertib (AZD1775). The cells that received sgRNAs were treated with IC25 and IC95 AZD1775 doses over 9 days (Fig.1a). Following successful quality control of the screen, which included confirming loss of representation of sgRNAs targeting essential genes (Supplementary Fig.1b), subsequent DrugZ^19^ bioinformatic analyses identified various factors predicted to contribute to WEE1 small-molecule inhibitor hypersensitivity or resistance. These include regulators of WEE1 stability, such as FAM122A^20^; factors downstream of WEE1, including CDK2 and CCNA2; components of nucleotide metabolism, RRM2 and DUT; and members of the anaphase-promoting complex, such as FZR1. We also noted that two genes involved in mRNA translation, GSPT1 and ALKBH8, emerged as hypersensitivity hits at the AZD1775 IC25 dose (Fig.1b). GSPT1 is an essential factor involved in translation termination that facilitates release of newly translated peptides from the ribosome via its GTPase domain^21^. ALKBH8 is a tRNA modifier that can modify 5- carboxymethyl uridine at the wobble position of the tRNA anticodon loop^22,23^.

**Fig. 1:**
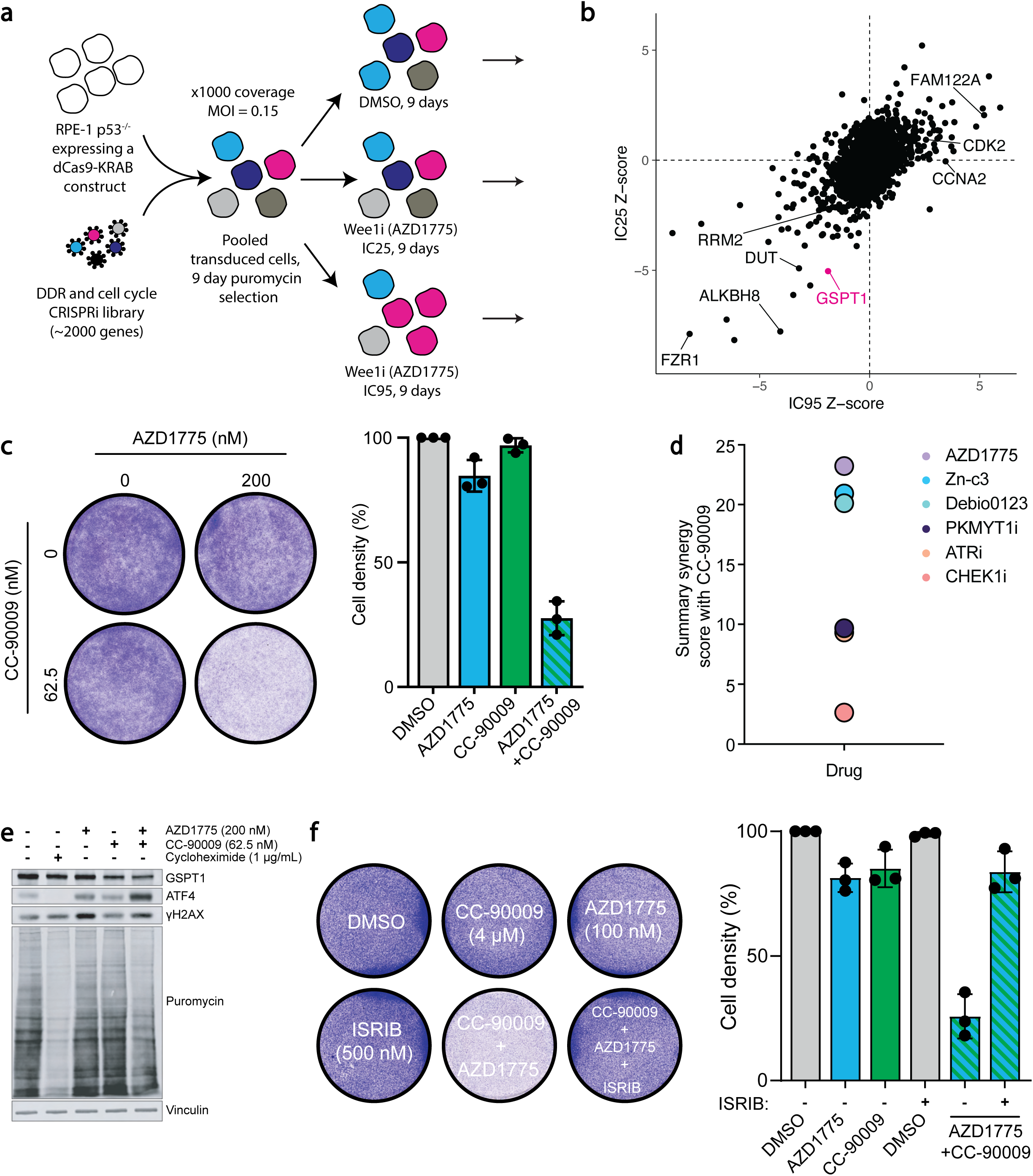
A pooled CRISPRi screen reveals the perturbation of GSPT1 as a sensitivity hit for WEE1 inhibitors. **a** Experimental design of pooled CRISPRi screen **b** CRISPRi screen results showing NormZ scores, calculated by DrugZ software, at IC95 (x-axis) and IC25 (y-axis) doses of AZD1775. Dosing information is available in Supplementary Fig.1a. **c** Representative image of 200 nM AZD1775 and 62.5 nM CC-90009 compounds alone or in combination for 72 hours in the RPE *TP53*^-/-^ cell line in a 6 well plate format followed by quantification of the cell density (biological n=3). Bar charts are depicted with means ± SD, points represent each biological replicate. **d** Resazurin cell viability summary synergy scores (combined Loewe, Bliss and HSA synergy scores) of CC-90009 in combination with WEE1 inhibitors (AZD1775, Zn-c3, Debio0123), PKMYT1i (RP-6306), ATRi (AZD6738) or CHEK1i (LY2603618) in RPE *TP53*^-/-^ cells in a 96 well plate format (biological n=4). Heatmaps of the drug combinations are shown in Supplementary Fig.4. **e** Western Blot showing 200 nM AZD1775 and 62.5 nM CC-90009 alone or in combination treated on the RPE *TP53*^-/-^ cell line for 24 hours. Cycloheximide (1 μg/mL) served as a positive control for global mRNA translation shutdown. Cells were treated with puromycin (5 μg/mL, 15 min) before harvesting. **f** Representative image of HAP1 cells treated with 100 nM AZD1775, 4 μM CC-90009 and 500 nM ISRIB compounds alone or in combination, for 72 hours in a 6 well plate format, followed by quantification of the cell density (biological n=3). Bar charts are depicted with means ± SD, points represent each biological replicate.

The above findings suggested a connection between WEE1 inhibitors and the control of mRNA translation. To explore this relationship, we employed a “molecular glue” compound that targets GSPT1 for degradation, CC-90009^24^. Thus, we found that CC-90009 was highly synergistic in combination with WEE1 inhibitors in both RPE-1 *TP53*^-/-^ and HAP1 cell lines as assayed via crystal violet staining (Fig.1c and Supplementary Fig.2, 3). Importantly, we found that three different small-molecule inhibitors of WEE1—AZD1775, Zn-c3, and Debio0123—exhibited strong synergy with CC-90009 in resazurin-based assays, which assess cell viability and metabolic activity. By calculating combined Loewe, Bliss and HSA synergy consensus scores^25^, we concluded that in RPE-1 *TP53*^-/-^ cells, WEE1 inhibitors were considerably more synergistic in combination with CC-90009 than were small-molecule inhibitors that targeted the WEE1-related kinase PKMYT1 (RP-6306^26^), or the DDR and replication stress kinases ATR (AZD6738^27^) and CHEK1 (LY2603618^28^) (Fig.1d and Supplementary Fig.4).

Perturbations of GSPT1 including CC-90009 treatment have been previously linked to ISR activation and the downstream accumulation of key transcription factors like ATF4^29,30^. Given this connection, we sought to investigate whether the synergy between WEE1 inhibitor and CC-90009 was dependent on ISR activation. A partial degradation of GSPT1 in combination with AZD1775 resulted in a substantial increase in ATF4 protein abundance (Fig.1e). In parallel, we also observed a synergistic reduction in global mRNA translation, as indicated by a decrease in puromycin incorporation into nascent protein, which serves as a measure of global cellular protein synthesis. Cycloheximide treatment, used as a positive control for mRNA translation shutdown, resulted in the expected reduction of puromycin incorporation^31^. These findings suggested that the ISR is implicated in the WEE1i– CC-90009 synergy. Indeed, we found that the synergistic loss of cell viability from combination of WEE1 inhibitor and CC-90009 could be rescued by inhibiting the integrated stress response by ISRIB^32^ or inhibiting GCN2 kinase activity by the compound GCN2iB^33^ in RPE-1 *TP53*^-/-^ and HAP1 cell lines (Fig.1f, and Supplementary Fig.2, 3). Furthermore, the loss of cell viability from treatments of AZD1775 or CC-90009 alone could also be rescued by ISRIB or GCN2iB compounds, which suggested that WEE1i–CC-90009 synergy arises from both drugs independently activating the ISR and GCN2 pathways (Supplementary Fig.2, 3).

### WEE1 small-molecule inhibitor treatment activates the ISR

Phosphorylation of the translation initiation factor eIF2α on Ser-51 to initiate the ISR is a vital signalling event that results in attenuating bulk protein synthesis as well as reprograming gene expression. This is mediated by the action of eIF2a kinases, including GCN2. GCN1, another key player in this pathway, can facilitate the activation of GCN2 under stress conditions^34,35^. A dose escalation of AZD1775 in the RPE-1 *TP53*^-/-^ cell line induced well-characterised markers of the ISR including phosphorylation of GCN2 at Thr-899, a known autophosphorylation site that correlates with GCN2 activation^36,37^, phosphorylation of eIF2α at Ser-51, and increased ATF4 protein levels. Inhibition of the GCN2 kinase by using 1 μM GCN2iB effectively blocked these markers of ISR induction (Fig.2a). Accordingly, we found that depletion of either GCN2 or GCN1 by CRISPRi induced resistance to AZD1775 in resazurin assays (Fig.2b, c). Furthermore, depletion of GCN1 in HEK293 cells significantly reduced the abundance of endogenous ATF4 protein induced by AZD1775 and, in a separate experiment, decreased the expression of a transfected ATF4 reporter compared to wild type, GCN1 expressing HEK293 cells (Supplementary Fig.5). Demonstrating that these effects were not limited to the previously mentioned cell lines, we observed that AZD1775 treatment increased nuclear ATF4 protein abundance across a panel of cancer and non-cancer cell lines; with such increases being abrogated upon co-treatment with 1 μM GCN2iB (Fig.2d).

**Fig. 2:**
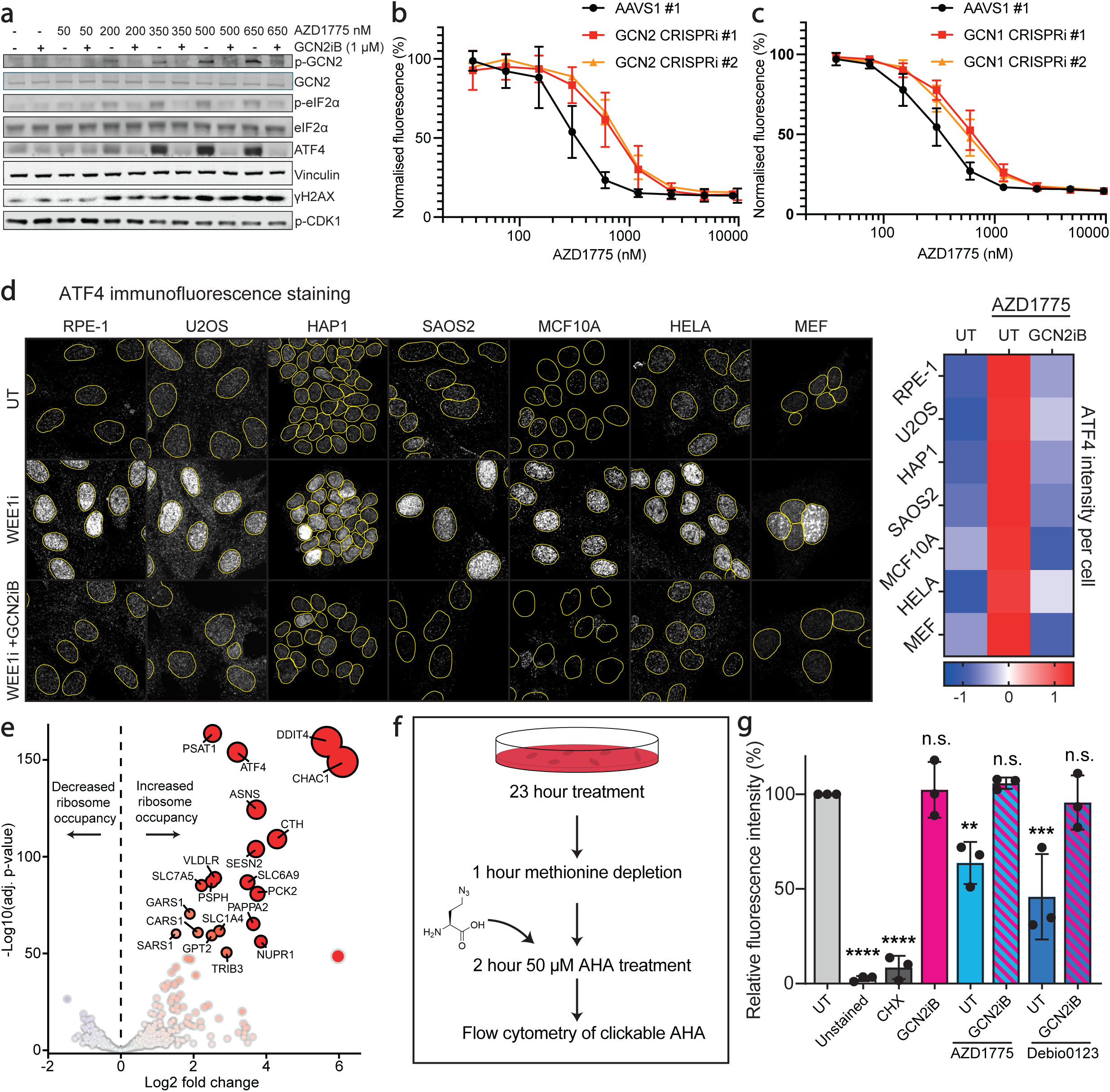
WEE1 small-molecule inhibitor treatment activates the ISR. **a** Western blot of RPE *TP53*^-/-^ cells treated for 24 hour treatments with increasing concentrations of AZD1775, with and without 1 μM GCN2iB. **b,c** Resazurin-based cell viability assay of RPE-1 *TP53*^-/-^ dCas9-KRAB cells expressing sgRNAs targeting GCN2, GCN1, or the AAVS1 locus. Cells were treated with varying concentrations of AZD1775 for 72 hours in a 96-well plate format (biological n=4). Graphs are depicted with means ± SD. Validations of the CRISPRi knockdown of GCN2 and GCN1 are shown in Supplementary Fig.9. **d** Representative immunofluorescence images of a panel of cell lines (RPE-1, U2OS, HAP1, SAOS2, MCF10A, HELA and MEF cell lines) probed for nuclear ATF4 treated with either DMSO, AZD1775 (650 nM for all cell lines except HAP1, which were treated at 300 nM), or AZD1775 in combination with 1 μM GCN2iB followed by row Z-score heatmap normalised per cell line summarising the immunofluorescence nuclear ATF4 intensity across different cell lines (biological n=4). **e** Ribosome profiling showing the log fold change of RPE-1 *TP53*^-/-^ cells treated with 650 nM AZD1775 vs DMSO for 10 hours (biological n=3). **f** Summary of flow cytometry based AHA experiment on the RPE-1 *TP53*^-/-^ cell line. An example of the flow cytometry gating strategy is available in Supplementary Fig.7a. **g** Bar chart of flow cytometry results showing the median fluorescence intensity of clickable AHA. RPE- 1 *TP53*^-/-^ cells (6 well plate format) were treated with 1 μg/mL cycloheximide, 350 nM AZD1775, 1.5 μM Debio0123 and 1 μM GCN2iB alone or in combination. A cell population not treated with AHA served as an unstained negative control. Median fluorescence intensity results were normalised to the untreated control (UT) (biological n=3). Bar charts are depicted with means ± SD, points represent each biological replicate. Statistical analyses were performed by one-way ANOVA with multiple comparisons, comparing to untreated condition, ns= not significant, ** *p*LJ<LJ0.01, *** *p*LJ<LJ0.001, **** *p*LJ<LJ0.0001.

Ribosome profiling, a technique that sequences RNA fragments that are protected by ribosomes^38^, revealed that AZD1775 treatment reprogrammed mRNA translation, increasing ribosome occupancy on several ISR-related transcripts, including ATF4 and TRIB3, as well as downstream effectors of ATF4, such as CHAC1 and PSAT1. Furthermore, regulators of the mTOR pathway, including DDIT4 and SLC7A5, also showed increased ribosome occupancy in response to AZD1775 treatment (Fig.2e), coinciding with previous observations that link mTOR pathway perturbations to WEE1 inhibitor resistance^39,40,41^.

To investigate whether the WEE1 inhibitors AZD1775 or Debio0123 affected global protein synthesis, we used L-AHA, a non-toxic methionine analogue that is incorporated into newly synthesised proteins in mammalian cells^42,43,44^. By measuring L-AHA incorporation, we assessed the rate of global nascent protein synthesis following treatment. We observed that both AZD1775 and Debio0123 reduced L-AHA incorporation, indicating a decrease in protein synthesis. Importantly, this reduction was rescued by co-treatment with 1 μM GCN2iB, suggesting that GCN2 activation is a key mediator of this effect (Fig.2f, g). The ribosome profiling data revealed no significant changes in codon occupancy at the P or A sites of the ribosome following AZD1775 treatment, suggesting that the observed reduction in nascent protein synthesis was not due to specific codon stalling (Supplementary Fig.6).

### WEE1 inhibitors synergise via ISR-dependent and -independent mechanisms

To determine which aspects of WEE1 inhibitor treatment are involved in ISR activation, we employed CRISPRi-based two-colour cell growth competition assays^45^. By utilising two distinct fluorescently labelled cell populations, this approach enabled us to evaluate the impact of different CRISPRi-mediated gene product depletions on cell fitness over time under WEE1 inhibitor treatment, both with and without ISR inhibition (Fig.3a). GFP/mCherry fitness graphs were generated over 9-day treatments from the two-colour cell growth competition assays. The area under the curve (AUC) of the GFP/mCherry fitness graphs was compared across different drug treatments, including AZD1775 both alone or in combination with ISRIB, or GCN2iB (Fig.3b).

**Fig. 3:**
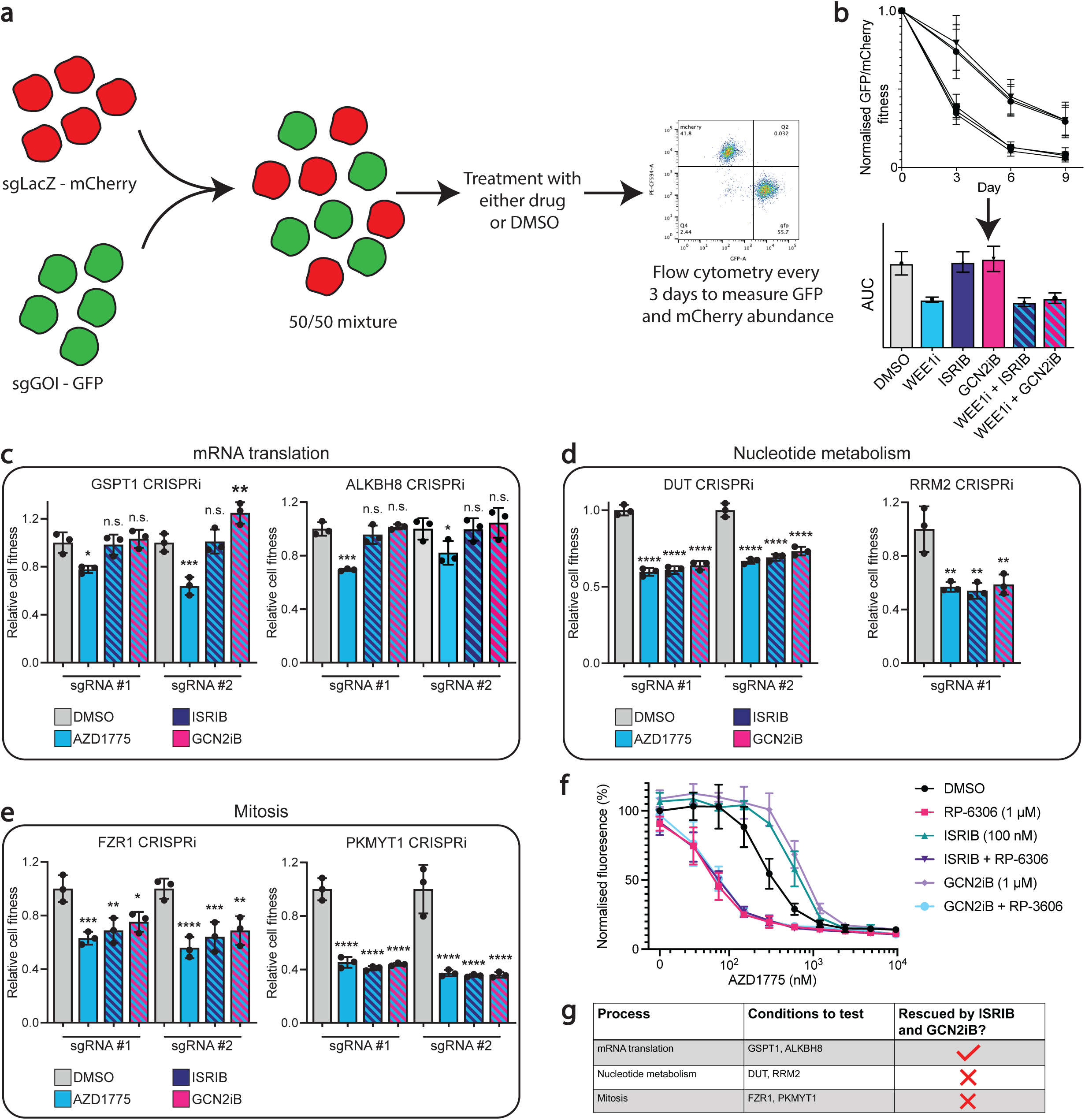
WEE1 inhibitors synergise via ISR-dependant and ISR-independent mechanisms. **a** Schematic showing flow cytometry based CRISPRi two-colour growth competition assays in RPE-1 *TP53*^-/-^ dCas9-KRAB cells. Cells were transduced with either sgLacZ-mCherry virus or sgGOI (gene of interest)-GFP. The mCherry and GFP expressing cell populations were mixed at a 50:50 ratio and treated with DMSO, AZD1775 (150 nM, 250 nM or 300 nM), 100 nM ISRIB or 1 μM GCN2iB, alone or in combination, for 9 days. Cells were assessed by flow cytometry, passaged and treated with fresh DMSO or drug every 3 days. The flow cytometry gating strategy is available in Supplementary Fig.7b. **b** Example of a normalised GFP/mCherry fitness graph. Values above 1 indicate the cell population expressing sgGOI causes increased relative cell fitness compared to cells expressing sgLacZ control, whereas values below 1 indicate decreased fitness. This is followed by an area under the curve that is generated from the GFP/mCherry fitness graph. **c, d, e** Bar charts quantifying the area under the curve of the normalised GFP/mCherry fitness graphs for different sgGOI-GFP populations vs sgLacZ-mCherry controls. Data were normalised to the respective DMSO-treated conditions. Values >1 indicate increased relative fitness compared to DMSO. Values <1 indicate decreased relative fitness compared to DMSO. Gray bars represent DMSO only, blue bars represent AZD1775 treatment only, blue/navy striped bars represent AZD1775 + 100 nM ISRIB, blue/pink striped bars represent AZD1775 + 1 μM GCN2iB. AZD1775 concentrations used: 150 nM for GSPT1, ALKBH8, and PKMYT1 CRISPRi; 250 nM for RRM2 and FZR1 CRISPRi; 300 nM for DUT CRISPRi. Original GFP/mCherry fitness graphs for all conditions (including sgAAVS1-GFP vs sgLacZ-mCherry controls) as well as ISRIB and GCN2iB only treatment controls are provided in Supplementary Fig.10-13, with CRISPRi knockdown validations in Supplementary Fig.9. Bar charts are depicted with means ± SD, points represent each biological replicate (biological n=3). Statistical analyses were performed by one-way ANOVA with multiple comparisons, comparing to DMSO treatment, ns= not significant, * *p*LJ<LJ0.05, ** *p*LJ<LJ0.01, *** *p*LJ<LJ0.001, **** *p*LJ<LJ0.0001. **f** Resazurin-based cell viability assay in RPE *TP53*^-/-^ cells treated with AZD1775 (varying concentrations) for 72 hours, with or without: DMSO, 1 μM PKMYT1i (RP-6306), 100 nM ISRIB, 100 nM ISRIB + 1 μM PKMYT1i, 1 μM GCN1iB and 1 μM GCN1iB + 1 μM PKMYT1i (biological n=3). Graphs are depicted with means ± SD. **g** Summary table of the different CRISPRi backgrounds tested.

CRISPRi-depleted cell populations that synergised with AZD1775, targeting gene products implicated in mRNA translation, nucleotide metabolism and mitosis, were tested. The gene products tested were various components identified as hits in the CRISPRi screen as previously discussed, with the exception of PKMYT1, which was not present in the CRISPRi library but is a known synergistic interactor of WEE1 perturbations^46,47^. The relative loss of cell fitness upon WEE1 inhibitor treatment in GSPT1- and ALKBH8-depleted populations was rescued by co-treatment with either ISRIB or GCN2iB, suggesting that the synergy between WEE1 inhibitor and mRNA translation defects is ISR- and GCN2-dependent (Fig.3c). By contrast, co-treatment with ISRIB or GCN2iB failed to significantly rescue the relative loss of cell fitness in DUT-, RRM2-, PKMYT1-, and FZR1-depleted populations upon WEE1 inhibitor treatment (Fig.3d, e). These findings strongly suggest that the synergy between WEE1 inhibitors and defects in nucleotide metabolism and mitosis is independent of the ISR and GCN2 kinase activity.

A recent study has reported that WEE1 inhibitors exhibit strong synergy with the PKMYT1 inhibitor RP-6306^46^; a combination currently being evaluated for safety and efficacy in phase 1 clinical trials (ClinicalTrials.gov: NCT04855656). Using resazurin viability assays, we found that the synergy of AZD1775 and RP-6306 remained unaffected by co-treatment with ISRIB or GCN2iB (Fig.3f). Given that WEE1 inhibitor and PKMYT1i synergy arises from the dysregulation of their shared phosphorylation target, CDK1, this suggests that CDK1 overactivation is not a driver of the WEE1 inhibitor-induced ISR phenotype. We also showed through immunofluorescence and western blotting analyses that depletion of other factors functioning downstream of WEE1, including CDK2, CCNE1, CCNE2 and CCNA2, did not impact AZD1775- induced ISR signalling (Supplementary Fig.8).

Furthermore, the synergistic relationship between AZD1775 and hydroxyurea^8^, which inhibits the ribonucleotide reductase enzyme^48^ to deplete nucleotide pools, was not impacted by ISRIB or GCN2iB co-treatments, as demonstrated by resazurin viability assays (Supplementary Fig.12b). This further supported the notion that the synergistic interaction between WEE1 inhibitors and defects in nucleotide metabolism was independent of the ISR.

### WEE1 inhibitors activate the ISR independent of WEE1

To determine whether ISR activation following WEE1 inhibitor treatment was driven by inhibition of WEE1 itself or by off-target effects of the small-molecule compound, we used recently developed cereblon dependent molecular glues that degrade WEE1 without requiring an ATP-competitive mechanism^49^. We established that in the RPE *TP53*^-/-^ cell line, both 1 µM HRZ-1-057-1 and 1 µM HRZ-1-098-1 induced considerable degradation of WEE1 within 1 hour (Fig.4a). We next carried out sequential drug-addition studies, wherein after a 1-hour molecular glue pre-treatment, cells were co-treated with DMSO or AZD1775 for 6 hours, and ISR signalling was assessed (Fig.4b). Notably, as revealed by western blotting, neither HRZ-1-057-1 nor HRZ-1-098-1 evoked ISR signals themselves; however, we continued to observe strong p-GCN2, p-eIF2α and ATF4 signals upon AZD1775 treatment, even after substantial degradation of WEE1 (Fig.4c). A similar phenomenon was observed with the ATP-competitive small molecule WEE1 inhibitors Zn-c3 and Derbio0123, both currently in clinical trials, and the experimental compound WEE1-IN-4, which also activated the ISR despite significant WEE1 degradation (Supplementary Fig.14a). From these data, we concluded that the ISR activation upon WEE1 inhibitor treatment was independent of WEE1 and due to off-target activity.

**Fig. 4:**
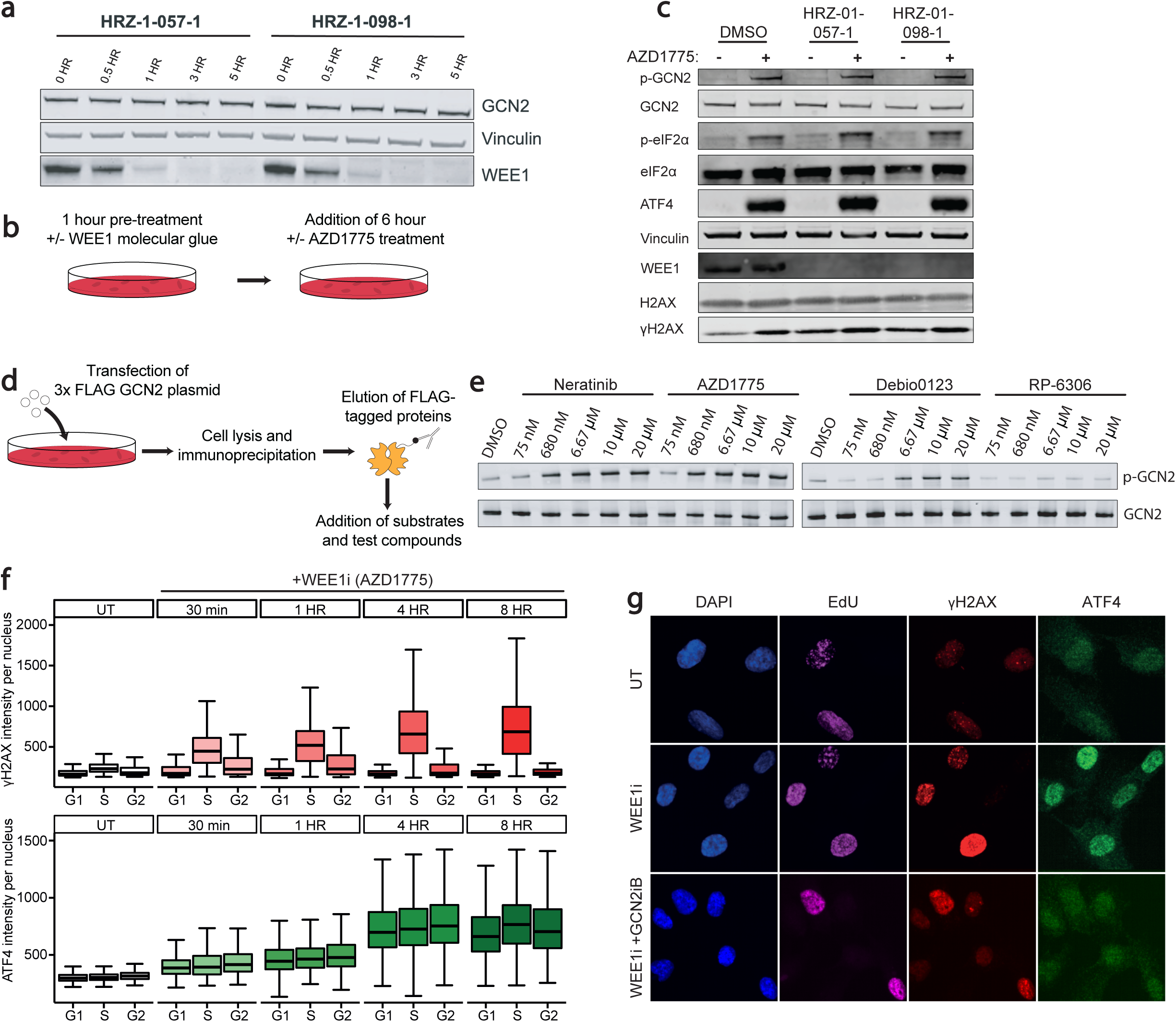
WEE1 inhibitors activate the integrated stress response independent of WEE1 and independent of cell cycle status. **a** Western blot showing the time course of WEE1 degradation in the RPE *TP53*^-/-^ cell line following treatment with 1 μM HRZ-057-1 or HRZ-1-098-1. **b** A schematic of the experiment in **(c)**. **c** Western blot of the RPE *TP53*^-/-^ cell line pre-treated with DMSO or 1 μM WEE1 molecular glues (HRZ-057-1 and HRZ-1-098-1) for 1 hour (total treatment duration: 7 hours), followed by DMSO or 650 nM AZD1775 for an additional 6 hours. **d** A schematic of the in vitro FLAG-tagged GCN2 experiment **e** Western blot of an in vitro experiment probing the total and phosphorylated GCN2 in the presence of DMSO, Neratinib, WEE1i (AZD1775 and Debio0123) and PKMYT1i (RP-6306). **f** Immunofluorescence analysis of nuclear ATF4 and γH2AX in the RPE *TP53*^-/-^ cell line treated with 650 nM AZD1775 across multiple timepoints (biological n=3). Box plots show median and quartile ranges. **g** Representative images showing ATF4 and γH2AX intensity in untreated (UT) cells and those treated with 650 nM AZD1775 alone or in combination with 1 μM GCN2iB for 24 hours in the RPE *TP53*^-/-^ cell line.

These observations raised the question of how the off-target activity was mediated. GCN2 has been recently reported to be activated by several ATP-competitive kinase inhibitors^50,36^, prompting us to hypothesise that WEE1 inhibitors might activate the ISR via a similar mechanism. To test this, we conducted *in vitro* phosphorylation assays with GCN2 in the presence of ATP and various small-molecule inhibitors. Thus, we observed that phosphorylation of GCN2 induced by AZD1775 was comparable to that of the known GCN2 activator, and EGFR inhibitor, Neratinib^36^. The Debio0123 compound was less potent at inducing GCN2 phosphorylation compared to Neratinib and AZD1775, but was still considerably more active in this regard than the PKMYT1 inhibitor, RP-6306 (Fig.4d, e and Supplementary Fig.14b). Notably, a previous *in vitro* kinome profile study had shown that high concentrations of AZD1775 bind to the second domain of GCN2 along with WEE1, WEE2, PLK1, among others^51^ (Supplementary Fig.14c). From this, we concluded that the off-target activity of AZD1775 and other WEE1 inhibitors was likely due to direct binding and activation of GCN2.

Additionally, we found that off-target ISR activation induced by AZD1775 was independent of cell cycle status. Thus, immunofluorescent staining revealed no significant differences in ATF4 signals across the G1, S and G2 phases of the cell cycle following AZD1775 treatment, whereas the DDR activation marker γH2AX (phosphorylated Ser-139 histone H2AX) was observed most strongly in S phase, as expected (Fig.4f, g). Furthermore, we observe that both ATF4 and γH2AX signals were induced very rapidly–within 30 minutes of AZD1775 treatment. Importantly, co-treatment with GCN2iB rescued ATF4 intensity but had no effect on γH2AX induction (Supplementary Fig.15), further emphasising that the DNA damage and ISR activation induced by AZD1775 treatment are independent toxicities.

### WEE1 PROTAC elicits less ISR toxicity compared to AZD1775

Given that AZD1775 activates the ISR, we were interested how this compound would compare to its PROTAC derivatives that use AZD1775 as a targeting warhead to degrade WEE1 (Fig.5a). Thus, we focused on ZNL-02-096, a previously developed WEE1 PROTAC that utilises AZD1775 as its warhead^49^. While AZD1775 exhibited strong synergy with CC-90009, ZNL-02-096 showed minimal synergy in RPE *TP53*^-/-^ cells (Fig.5b and Supplementary Fig.16a). The depletion of GCN2 by CRISPRi gave rise to the resistance of AZD1775 as previously shown in Fig.2b but not of ZNL-02-096 (Fig.5c).

**Fig. 5:**
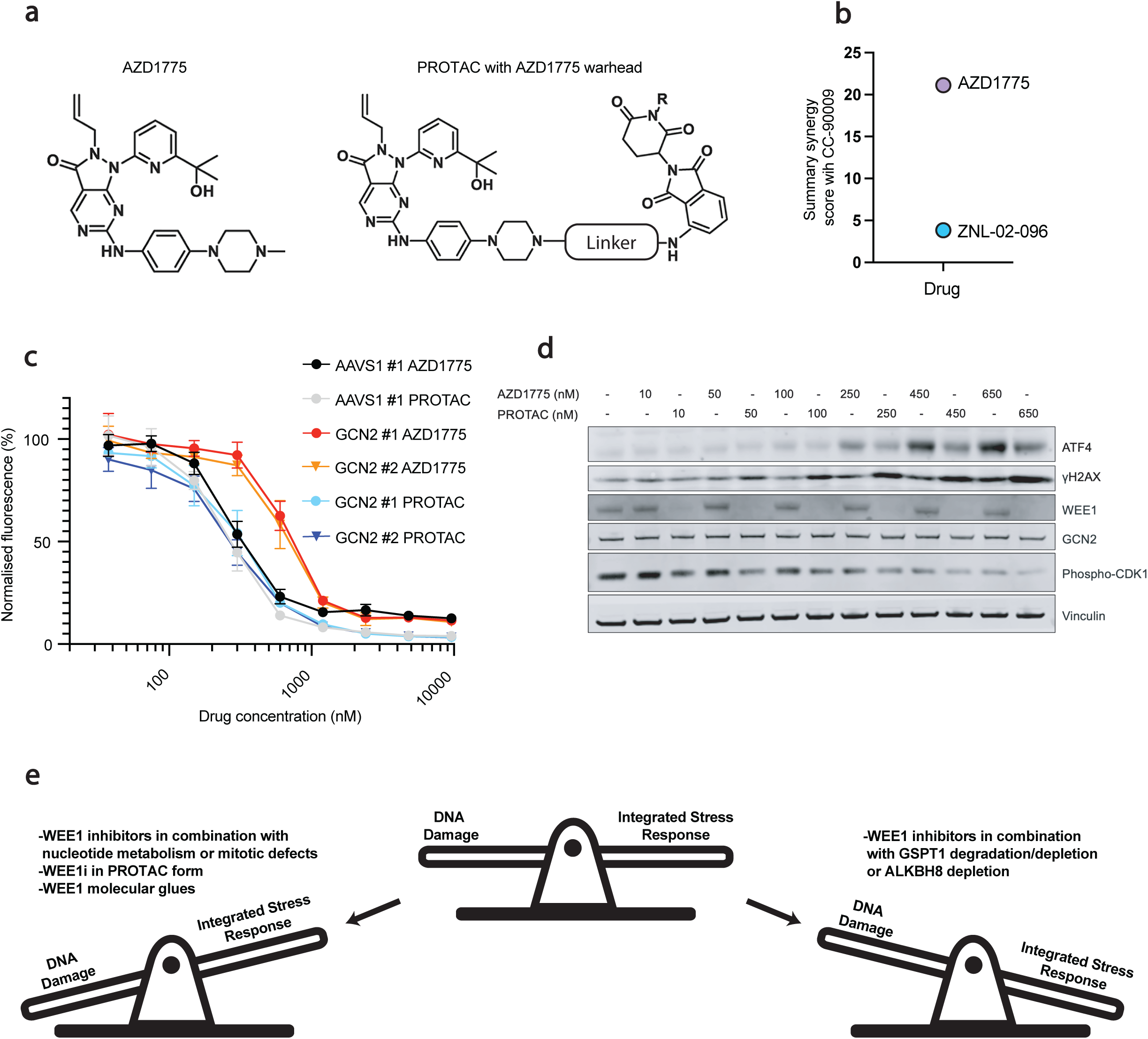
WEE1 PROTAC elicits less ISR toxicity compared to AZD1775. **a** Chemical structure of AZD1775 and PROTACs utilising AZD1775 as a warhead. **b** Resazurin cell viability summary synergy scores (combined Loewe, Bliss and HSA synergy scores) of CC-90009 in combination with AZD1775 or ZNL-02-096 in the RPE *TP53*^-/-^ cell line in a 96 well plate format (biological n=3). Heatmaps of the drug combinations can be found in Supplementary Fig.16. **c** Resazurin cell viability assay with varying concentrations of AZD1775 or ZNL-02-096 treated for 72 hours in a 96 well plate format in the RPE-1 *TP53*^-/-^ dCas9-KRAB cell line expressing sgRNAs that target GCN2 or the AAVS1 locus (biological n=3). Graphs are depicted with means ± SD. **d** Western blot comparing AZD1775 and ZNL-02-096 18-hour treatments in the RPE *TP53*^-/-^ cell line. **e** Schematic showing the balance between the two independent toxicities of DNA damage and ISR activation for WEE1 inhibitor treatments.

The above data suggested that AZD1775 in its PROTAC form had less ISR mediated toxicity. Indeed, after 18 hours of treatment of the RPE *TP53*^-/-^ cell line, ZNL-02-096 induced less ATF4 protein abundance than AZD1775. However, despite eliciting a weaker ISR response, ZNL-02-096 induced greater γH2AX levels than an equivalent molar concentration of AZD1775 (Fig.5d). Comparing across multiple timepoints, we observed that ZNL-02-096 treatment did give rise to ISR signals comparable to those elicited by AZD1775; however, with ZNL-02-096, these signals were transient and dissipated more rapidly over time. We also tested another WEE1 PROTAC, ZNL-02-047 that like ZNL-02-096, employs AZD1775 as a warhead but differs in linker composition. Whereas ZNL-02-096 employs a hydrocarbon linker, ZNL-02-047 features a polyethylene glycol linker^52^. Notably, ZNL-02-047 induced considerably weaker ISR activation compared to both AZD1775 and ZNL-02-096 at the same 650 nM concentration (Supplementary Fig.16b). Taken together, these findings suggested that PROTAC derivatives of AZD1775 may serve as a suitable alternative to AZD1775 if the desired outcome is to induce genotoxicity whilst limiting ISR activation.

## DISCUSSION

We have investigated the determinants of sensitivity of RPE-1 *TP53*^-/-^ cells to the ATP competitive WEE1 kinase inhibitor AZD1775. Our findings revealed that this WEE1 inhibitor synergised with depletion of mRNA translation factors GSPT1 and ALKBH8 through activation of the ISR and the GCN2 kinase. Furthermore, we established that treatment with AZD1775, as well as other WEE1 inhibitors— including Zn-c3, Debio0123, and WEE-IN-4—activated the ISR via GCN2. This aligns with a recent study reporting ISR activation following AZD1775 treatment in small-cell lung cancer (SCLC) cell lines^53^. Importantly, we have shown that this phenotype is conserved across cancer, non-cancer, human and mouse cell lines; and additionally, that this WEE1 inhibitor-induced ISR activation is a ‘blanket’ toxicity that is independent of cell cycle status.

Prolonged ISR activation can lead to cell death^54^, and it is well documented that DNA damage can also give rise to cell death^55^. Therefore, we suggest that currently exploited WEE1 inhibitors should regarded as inhibitors that can elicit a dual toxicity—both ISR activation and genotoxicity. Through CRISPRi-based cell growth competition assays, we have demonstrated that hypersensitivity to WEE1 inhibitors can be exploited via both ISR-dependent and ISR-independent mechanisms. Additionally, we also showed this with different drug combinations. Thus, we found that CC-90009, a GSPT1 degrader, exhibited strong synergy with WEE1 inhibitors through ISR activation, whereas the combination of WEE1 inhibitors with PKMYT1 inhibitors or hydroxyurea displayed potent synergy in an ISR-independent manner.

Notably, we found that recently developed WEE1 molecular glues that target WEE1 outside of the ATP-binding pocket showed no evidence of ISR activation but effectively induced on-target genotoxicity. By degrading WEE1 using molecular glues, followed by treatment with WEE1 small-molecule inhibitors, we demonstrated that these small-molecule inhibitors activated the ISR independent of the WEE1 kinase. Furthermore, our biochemical data strongly suggested that WEE1 small-molecule inhibitors directly target the GCN2 kinase. However, it remains unclear why no GCN2 degradation was observed with the WEE1 PROTACs ZNL-02-096 and ZNL-02-047. Notably, depletion of both GCN1 and GCN2 rescued AZD1775-induced toxicity, and GCN1 knockout cells showed greatly reduced ATF4 ISR signalling in response to AZD1775 and low-doses of GCN2iB (Supplementary Fig.5; GCN2iB has been noted to activate GCN2 at low doses^56^). One possible explanation for the importance of GCN1 is that it ‘primes’ GCN2 for activation by certain small-molecule inhibitors, potentially by increasing ATP-binding pocket accessibility. This hypothesis may also explain why WEE1 inhibitors synergise with mRNA translation defects, as these conditions could promote increased GCN1-GCN2 interactions.

In conclusion, we have established that various WEE1 small-molecule inhibitors activate the ISR via GCN1 and GCN2 in an off-target manner across a wide range of cell lines. We demonstrated that AZD1775 can synergise with various genetic backgrounds and drug treatments via both ISR-dependent and ISR-independent mechanisms (Fig.5e). Our study suggests that WEE1 small-molecule inhibitors used in the clinic lack precise specificity towards WEE1. However, using molecular glues and PROTACS that degrade WEE1 causes greatly reduced ISR toxicity while maintaining WEE1-dependent genotoxicity. Therefore, such modalities may represent more precise and effective therapeutic alternatives for targeting WEE1- mediated toxicity without ISR related side-effects.

## MATERIALS AND METHODS

### Methods

#### Cell culture

Cell lines used in this study were routinely tested for mycoplasma contamination with no contaminations found. U2OS, SAOS2, HeLa, MEF and HEK293 cells were cultured in Dulbecco’s modified Eagle’s medium (DMEM, Gibco A4192101), hTERT RPE-1 (WT, *TP53*^-/-^ and *TP53*^-/-^ dCas9-KRAB) in Dulbecco’s Modified Eagle Medium: Nutrient Mixture Ham’s F-12 (DMEM/F-12, Gibco 11320033), HAP1 cells in Iscove’s modified Dulbecco’s medium (Sigma I3390-500mL), and MCF10A cells in mammary epithelial cell growth medium supplemented with bovine pituitary extract, human epidermal growth factor, insulin, hydrocortisone and cholera toxin (MEGM, Lonza CC-3150). All cell lines were grown at 37 °C and 5% CO_2_. All cell lines except for MCF10A and RPE-1 (RPE-1 were cultured without additional L-glutamine supplementation) were cultured in media supplemented with 10% foetal bovine serum (FBS), 2 mM L-glutamine, 100 units/mL penicillin, and 100 μg/mL streptomycin.

hTERT RPE-1 *TP53*^-/-^ dCas9-KRAB cell line was provided by the Jacob Corn Lab and were generated by transduction with a lentiviral vector encoding the dCas9- KRAB and a Blasticidin resistance cassette with a low MOI. Cells were selected with Blasticidin, and single cell clones were seeded by cell sorting. Resulting clones were validated for CRISPRi activity by CD55 knockdown efficiency.

Methionine free medium was made as follows: 7.4 g of DMEM/F12 powder (D9785, Sigma), 0.077 g calcium chloride (Sigma), 0.046 g L-lysine (Sigma), 0.03 g L-leucine (Acros Organics), 0.0244 g magnesium sulfate (Fisher Scientific), 0.0306 g magnesium chloride (Sigma) and 0.6 g sodium bicarbonate (Sigma) was dissolved in 500 mL of MilliQ water and filtered using Millipore express PLUS 0.22 µm PES filter. 10% FBS and 100 units/mL penicillin and 100 µg/mL streptomycin was added.

#### Compounds

ATR inhibitor (AZD6738), CHEK1 inhibitor (LY2603618), WEE1 inhibitors (AZD1775, Zn-c3 and Debio0123), PKMYT1 inhibitor (RP-6306) and ISRIB were obtained from SelleckChem. GCN2iB and WEE1-IN-4 was obtained from Medchemexpress. WEE1 PROTAC (ZNL-02-096) was obtained from Tocris Bioscience. Hydroxyurea and cycloheximide was obtained from Sigma-Aldrich. WEE1 molecular glues (HRZ-1- 057-1, HRZ-1-098-1) and WEE1 PROTAC (ZNL-02-047) were kindly provided by the Nathanael S. Gray laboratory. In the original publication, HRZ-1-057-1 is referred to as ’compound 1,’ and HRZ-1-098-1 is referred to as ’compound 10’^49^.

#### Cell line generation for CRISPRi cell lines

To generate CRISPRi cell lines, lentiviruses were first generated in LentiX 293T cells by transfecting packaging plasmids psPAX2 (Addgene, #12260) and pMD2.G (Addgene, #12259) with the plasmid of interest using the transfection reagent TransIT-LT1 (Mirus Bio) according to the manufacturer’s protocol. 72 h later, medium was collected, centrifuged at 2000g for 10 minutes and the lentivirus containing supernatant was stored at −80D°C. The RPE-1 *TP53*^-/-^ dCas9-KRAB were cultured in 2 μg/mL puromycin 1 day post-transduction to select cells that had incorporated an sgRNA. All CRISPRi cell lines tested in the study were validated either by western blot or by RT-qPCR. sgRNA protospacers and primers used can be found in Supplementary Table 2.

#### sgRNA cloning

CRISPRi sgRNAs were cloned into either mCherry- or GFP-containing lentiviral vectors (addgene #185473 or addgene #185474 plasmids) Forward and reverse primers for each sgRNA were annealed by pre-incubation at 37°C for 30 minutes with T4 polynucleotide kinase (PNK; NEB), followed by incubation at 95°C for 5 minutes and then ramp down to 25°C at 5°C/min. Annealed sgRNAs were ligated into the corresponding vector that had been digested with BsmBI restriction enzymes using T4 Ligase (NEB). All sgRNA protospacers are listed in Supplementary Table 2.

#### CRISPRi library

For the DDR and cell cycle CRISPRi library design, the two strongest sgRNAs against each gene were selected from the CRISPRi-v2 library (PMID: 27661255). Where possible, this was determined using empirical scores, otherwise the predicted activities from the CRISPRi v2 library design algorithm was used. In case of multiple likely transcripts, sgRNAs against all transcripts in the CRISPRi-v2 library were included, such that some genes are targeted by 4 or more (up to 8) perfectly matched sgRNAs. sgRNA sequences of the library can be found in Supplementary Table 3. Protospacer sequences were appended with BlpI and BstXI restriction sites and PCR adapter. Oligonucleotides were synthesised by Agilent Technologies and cloned into the pLG1 library vector (pU6-sgRNA Ef1alpha-Puro-T2A-BFP). To ensure accurate representation of the library, Next-Generation Sequencing using the Illumina MiSeq platform was performed.

#### CRISPRi screen

RPE-1-hTERT dCas9-KRAB *TP53*^-/-^ cells were transduced with a lentiviral CRISPRi library (DDR CRISPRi allelic series library targeting 2294 genes with pLG1 (pCRISPRia- v2) backbone provided by the Corn Lab, ETH Zurich) at an MOI of 0.15 via spinfection. Two separate biological replicates of the screen were performed i.e., two separate library transductions. The next day following transduction, fresh medium containing puromycin (2 μg/ml) was added. Cells were negatively selected with puromycin for a total of 9 days, at which point each replicate was divided into different treatments and subcultured every 3 days for a total of 9 days. Cell pellets were frozen at the end of the day 9 treatment (day 19 post-transduction) and as well as day 3 post-transduction for gDNA isolation. Library coverage of at least x750 cells per sgRNA was maintained at every step for the DMSO and the IC25 arm, and at least x300 cells per sgRNA for the IC95 arm. gDNA from cell pellets were isolated using the QIAamp Blood Midi Kit (Qiagen, Cat# 51185) and genome-integrated sgRNA sequences were amplified by PCR using NEBNext Ultra II Q5 Mastermix (New England Biolabs, Cat# M5044L). Sample preparations were performed by amplifying the sgRNA with forward and reverse primers containing single TruSeq indexes. The final magnetic bead-purified products (purified using SPRI MBSpure beads obtained from the Vienna Biocenter, Austria) were sequenced on an Illumina HiSeq3000 system to determine sgRNA representation in each sample. A 15% PhiX spike-in was used. To identify both synergistic and suppressor interactions, sgRNAs enriched or depleted in the treated samples were determined by comparison to control samples using DrugZ software^19^. Exorcise software was used to verify targeting of the sgRNAs to genome assembly GRCh38^57^.

#### Immunoblotting

Cells were collected in lysis buffer (50 mM Tris-HCl, pH 6.8, 1% SDS, 5 mM EDTA) and heated for 8 min at 95 °C. Protein concentrations were determined using a NanoDrop spectrophotometer (Thermo Scientific NanoDrop One) at 280 nm. NuPage x4 LDS loading buffer (Invitrogen, NP0007) with 100 mM DTT was added to protein lysates at a final concentration of x1. SDS–PAGE was performed to resolve proteins on pre-cast NuPAGE Novex 4–12% Bis/Tris gradient gels (Invitrogen).

Separated proteins were transferred to nitrocellulose (Sigma GE10600002), blocked for 1 hour at room temperature in 5% BSA 0.1% tween 20 TBST and immunoblotted with the indicated primary antibodies overnight at 4°C. The nitrocellulose membranes were washed 3 times for 10-minute washes in 0.1% TBST. The membranes were incubated in 1:10,000 secondary antibody for 1 hour at room temperature and then washed 3 times for 10-minute washes in 0.1% TBST. All secondary antibodies (except for probing ATF4) were incubated in fluorescent-conjugated secondary antibody (anti-rabbit secondary: LI-COR 926-68021, anti-mouse secondary: LI-COR: 926-32212). HRP-conjugated secondary antibody (Santa-Cruz, sc-2313) was used to probe for ATF4. Chemiluminescent signal was detected using SuperSignal West Pico PLUS (Thermo Scientific, 34580). Western blotting images were captured using ChemiDoc MP Imaging System (Bio-Rad) or LI-COR Odyssey M. A list of primary antibodies and dilutions used in this study can be found in Supplementary Table 1.

#### Crystal violet staining

Cells were added directly to drugs in a 6 well plate format with a total volume of 2 mL medium per well. The RPE *TP53*^-/-^ cell line was seeded at 50,000 and 5,000 cells per well for 3 day and 6 day treatments respectively. The HAP1 cell line was seeded for 250,000 and 25,000 cells per well for 3 day and 6 day treatments respectively. Cells were incubated with drug at 37°C and 5% CO_2_. Following 3 days or 6 days, the 6 well plates were washed with PBS and then stained with crystal violet (Pro-Lab PL.7002) for 30 minutes. Following this, the 6 well plates were washed with water, dried and imaged on an Epson Perfection V800 scanner. The cell density was quantified using ImageJ software.

#### Resazurin assays

Cells were added directly to drugs in a 96 well plate format with a total volume of 200 µL medium per well. RPE *TP53*^-/-^, RPE-1 *TP53*^-/-^ dCas9-KRAB and HAP1 cell lines were seeded at 750, 1000 and 6000 cells per well respectively. Cells were incubated with drug at 37°C and 5% CO_2_ for 72 hours. Following this, pre-warmed Resazurin reagent was added at a final resazurin concentration of 20 µg/mL and incubated for 3 hours. Resazurin stock reagent was prepared by dissolving resazurin powder (Thermo Fisher Scientific, 418900250) in sterile PBS at 200 µg/mL and filtered with a 0.22 µm filter. After 3 hours, the 96 well plates were read on the CLARIOstar plate reader with a 530-560 nm excitation wavelength and 590 nm emission wavelength.

200 µL medium only outer wells were used as negative controls to account for background fluorescence. The fluorescence values were normalised to the DMSO only treated wells. Synergy scores from the normalised fluorescence intensities was calculated using the SynergyFinder software (<u>synergyfinder.fimm.fi</u>)^25^.

#### CRISPRi-based two colour competitive growth assay

RPE-1 *TP53*^-/-^ dCas9-KRAB cells were infected at an MOI of ∼0.2 and treated with 2 µg/mL puromycin the next day and maintained in puromycin until the population was fully selected. This ensured that the transduced cell population all contained a puromycin resistance plasmid of NLS-mCherry LacZ-sgRNA or NLS-GFP GOI- sgRNA. Following selection, the mCherry- and GFP-expressing cells were mixed 1:1 (10,000 cells + 10,000 cells) and plated with or without drug in 12 well plate format. During the experiment, cells were subcultured and re-treated with or without drug every three days when they reached near-confluency. Cells were analysed on the A5 FACS Symphony (BD Biosciences), gating for GFP- and mCherry signal the day of the initial plating (t=0) and on days 3, 6 and 9. The efficiency of the knockdown by CRISPRi was analysed by western blotting or by RT-qPCR.

#### Immunofluorescence

Cells were seeded in 24 well glass bottom plates (Greiner 662892) at 50,000 cells per well and incubated for 24 hours. Cells were treated with either WEE1 inhibitor, or WEE1 inhibitor and GCN2 inhibitor for indicated times by exchanging medium with fresh medium containing drugs. Cells were washed once with PBS, fixed with 4% PFA for 10 minutes at room temperature, permeabilised with PBS +0.1% Tween-20 (PBST) +0.2% Triton X-100 for 20 minutes at room temperature and blocked with 5% BSA in PBST for 1 hour at room temperature. Antibodies were diluted to desired concentrations in 5% BSA PBST and incubated at 4°C overnight. Cells were wash four times in PBST for 5 minutes at room temperature. Secondary antibodies tagged with Alexa Fluor 488 (Thermo, A11029) or Alexa Fluor 568 (Thermo, A11036) were diluted 1:1000 in 5% BSA PBST +1 μg/mL DAPI and incubated for 1 hour at room temperature. Cells were wash four times in PBST for 5 minutes at room temperature and 2mL PBS wash added to each well for storage until imaging. For cell cycle stratification, 1 μM EdU was added to cell culture medium 30 minutes before fixing, and following secondary antibody a CLICK reaction was performed to conjugate Alexa Fluor 647 azide (Thermo, A10277) to the incorporated EdU as follows; 2mM copper sulphate, 1 μM Alexa Fluor 647 azide and 10mM sodium ascorbate. Following CLICK, cells were wash twice with PBST for 5 minutes at room temperature before storing. Plates were imaged using an Opera Phenix Plus (Revvity) high content spinning disk confocal microscope and analysis was completed in the Harmony software (Revvity). Nuclei were identified through DAPI signal and fluorescent intensities were quantified per nucleus as a mean intensity.

#### AHA incorporation assay

RPE-1 *TP53*^-/-^ cells were seeded in a 6 well plate with a total volume of 2 mL medium per well. Wells were treated with or without drug the next day for 23 hours. Following this, the 6 wells were washed with PBS and treated for 1 hour in 2 mL of methionine free cell medium that contained the same drug concentrations as previous for each well. 50 µM L-Azidohomoalanine/AHA (Thermo, C10102) was added for 2 hours following a 1 hour methionine depletion. The 6 well plates were washed with PBS, typsinised and harvested. Following fixation, click reactions using Click-iT Cell Reaction Buffer (Thermo, C10269) and 2.5 µM Alexa Fluor 647 Alkyne, (Thermo, A10278) and subsequent washes were performed as per manufacturer instructions using the flow cytometry experimental protocol for adherent cells. Cells were analysed and gated on the A5 FACS Symphony (BD Biosciences) and analysed with FlowJo v.10.8.1. An example of the flow cytometry gating is shown in Supplementary Fig.7a.

#### RT-qPCR

RNA was extracted from snap frozen cell pellets using the RNeasy mini kit (Qiagen) or Maxwell RSC simplyRNA Cells Kit (Promega). RNA extraction was as per manufacturer’s instructions. During the extraction protocol, RNase-free plastic ware and solutions were used. After RNA extraction, isolated RNA was quantified using a Nanodrop spectrograph, and stored at −80 °C. cDNA was reverse transcribed from 1 μg of RNA using SuperScript IV VILO Master Mix (Invitrogen, 11756050), according to the manufacturer’s instructions. cDNA was stored at −20 °C. qPCR was performed using 1 μL of cDNA,10 μL of 2× Fast SYBR Green Master mix (Thermo Fisher Scientific, , 4385612) and 500 nM forward and reverse primers, in a final volume of 20 μL. Primers were designed and ordered to span an exon-exon junction of the target genes (Sigma-Aldrich) and sequences are detailed in Supplementary Table 2. qPCR analysis was performed on an QuantStudio 5 Real-Time PCR Systems (Thermo Fisher Scientific), in technical triplicates. Gene expression changes were calculated using the 2–ΔΔCt method^58^. RT qPCR results can be found in Supplementary Fig.9b, c.

#### Ribosome profiling

RPE *TP53*^-/-^ cells were treated either with DMSO or 650 nM AZD1775 for 10 hours in 15 cm^2^ plates. Following the incubation, the growth media was aspirated and washed in ice-cold PBS/CHX (1 X PBS supplemented with 100 µg/mL cycloheximide). Following this, 3 mL of ice old PBS/CHX was added per dish, cells were scraped extensively and pelleted at 300G centrifugation. Supernatant was aspirated and the pellets were snap frozen. Flash frozen cell pellets were lysed in ice-cold polysome lysis buffer (20 mM Tris pH 7.5, 150 mM NaCl, 5 mM MgCl_2_, 1 mM DTT, 1% Triton X-100) supplemented with cycloheximide (100 µg/mL). For ribosome profiling (Ribo-seq), remaining lysates were digested in the presence of 35U RNase1 for 1 hour at room temperature. Following RNA purification, PNK end repair and size selection of ribosome protected mRNA fragments on 15% urea PAGE gels, contaminating rRNA was depleted from samples using EIRNABio’s custom biotinylated rRNA depletion oligos. Enriched fragments were converted into Illumina compatible cDNA libraries. Ribo-seq libraries were sequenced 150PE on Illumina’s Nova-seq 6000 platform to depths of 100 million raw read pairs per sample.

#### Flag tagged GCN2 in vitro assay

*EIF2AK4*^-/-^ HeLa cells were transfected with pcDNA3.1/hygro(-)_hGCN2-3xFLAG^59^ to express 3xFLAG-tagged GCN2. At 24 hours post transfection, cell lysates were prepared (50 mM Tris, pH 7.4, 150 mM NaCl, 1 mM EDTA, 1% Triton-X, protease and phosphatase inhibitors). FLAG tagged protein was purified using anti-FLAG® M2 affinity gel slurry (Millipore, A2220) and eluted with 25 μg/ml 3xFLAG peptide solution (Pierce™ 3x DYKDDDDK, Thermo Scientific, A36805). 10 μL of eluate were incubated with recombinant eIF2α-NTD, amino acids 2–187 (gifted by Heather Harding) to a final concentration of 1 μM and 500 μM ATP in reaction buffer: 50 mM HEPES, pH 7.4, 100 mM potassium acetate, 5 mM magnesium acetate, 250 μg/ml BSA, 10 mM magnesium chloride, 5 mM DTT, 5 mM β-glycerophosphate. Reactions were incubated at 32°C for 10 minutes, then immediately quenched with 5 μL of 6X SDS sample buffer for immunoblotting and heated for 8 minutes at 95°C.

#### GCN1 ATF4 reporter transfection

Wild-type and GCN1^-/-^ HEK293 cells (ab266780) were transfected with pGL4.2_CMV_hATF4UTR:nLucSTOP to express the hATF4::nanoLuc reporter^59^. In 384-well plates, 103 cells in 20 μL per well were added to 5 μL of 5X treatment solution in 10% FBS-DMEM, yielding final concentrations of 5, 10, 35, or 70 nM GCN2iB, 3 mM histidinol, and 50, 100 or 200 nM AZD1775. After 6 h of incubation at 37°C, 25 μL of NanoGlo® luciferase assay reagent were added to each well and bioluminescence (360-545 nm) acquired (Tecan Spark plate reader).

## ACKNOWLEDGEMENTS

We thank members of the Jackson laboratory, as well as members of the Marciniak, Villunger and Loizou laboratories for their assistance, discussion and reagents. We thank the Jacob Corn laboratory, ETH Zurich for their discussions and for providing the DDR focused CRISPRi library and RPE-1 dCas9-KRAB cell line. We are grateful to the Nathanael S. Gray laboratory, Stanford University for providing HRZ-1-057-1, HRZ-1-098-1 and ZNL-02-047 WEE1 degrader compounds. We also acknowledge the Research Instrumentation and Cell Services, Flow Cytometry and Microscopy Core Facilities at the CRUK Cambridge Institute for technical support and access to equipment and reagents. We thank the labs of Marcel Van Vugt (Groningen) and Daniel Durocher (Tornonto) for their discussions and sharing of unpublished data. We also thank Anna Schrempf (Loizou/Winter labs, Vienna) and Gonçalo Oliveira (Loizou/Hanzlíková labs, Vienna/Prague) for their discussions and sharing of CRISPR screening protocols. We thank Manu Hegde and Eszter Zavodszky, LMB, Cambridge, for their insightful discussions and assistance with ribosomal assays. We also appreciate the informative discussions with members of the AstraZeneca Oncology R&D team in Cambridge. CRISPRi libraries were sequenced by Biomedical Sequencing Facility (BSF) (https://www.biomedical-sequencing.org). Ribo-seq libraries were generated and sequenced by EIRNABio (https://eirnabio.com). This research was supported by the European Research Council (ERC) under the European Union’s Horizon 2020 research and innovation programme (Grant agreement No. 855741-DDREAMM-ERC-2019-SyG). Research in the SPJ laboratory is supported by Cancer Research UK (CRUK) Discovery Award DRCPGM\100005, CRUK core grant C9545/A29580 and SEBINT- 2024/100003 and ERC Synergy Award 855741 (DDREAMM); AH is funded by a grant to SPJ from GSK; CGG by an A*STAR National Science Scholarship; FBD by a Harding Distinguished Postgraduate Scholarship; SL by ERC Synergy Award 855741; and ASB by Wellcome Early Career Award (227014/Z/23/Z). SJM is supported by the MRC, MCMB MR/Y011813/1. JZ is funded by the Doctoral Training Programme in Medical Research (DTP-MR) and the Cambridge Trust. JEC is supported by the NOMIS Foundation, the Lotte and Adolf Hotz-Sprenger Stiftung, the Swiss National Science Foundation (project grants 310030_188858, 310030_201160 and 320030-227979), and the European Research Council (ERC) under the European Union’s Horizon 2020 research and innovation program (grant agreement No 855741, DDREAMM). JEC is a co-founder and SAB member of Serac Biosciences and an SAB member of Mission Therapeutics, Relation Therapeutics, Hornet Bio, and Kano Therapeutics. The lab of JEC has funded collaborations with Allogene, Cimeio, and Serac. JF is a recipient of the EMBO Postdoctoral Fellowship (ALTF 220-2021). MFS is a recipient of the Boehringer Ingelheim Fonds (BIF) PhD fellowship.

