## Supplemental Material (Supp Figures, Supp Table 1 & 2) for "WEE1 inhibitors synergise with mRNA translation defects via activation of the kinase GCN2"

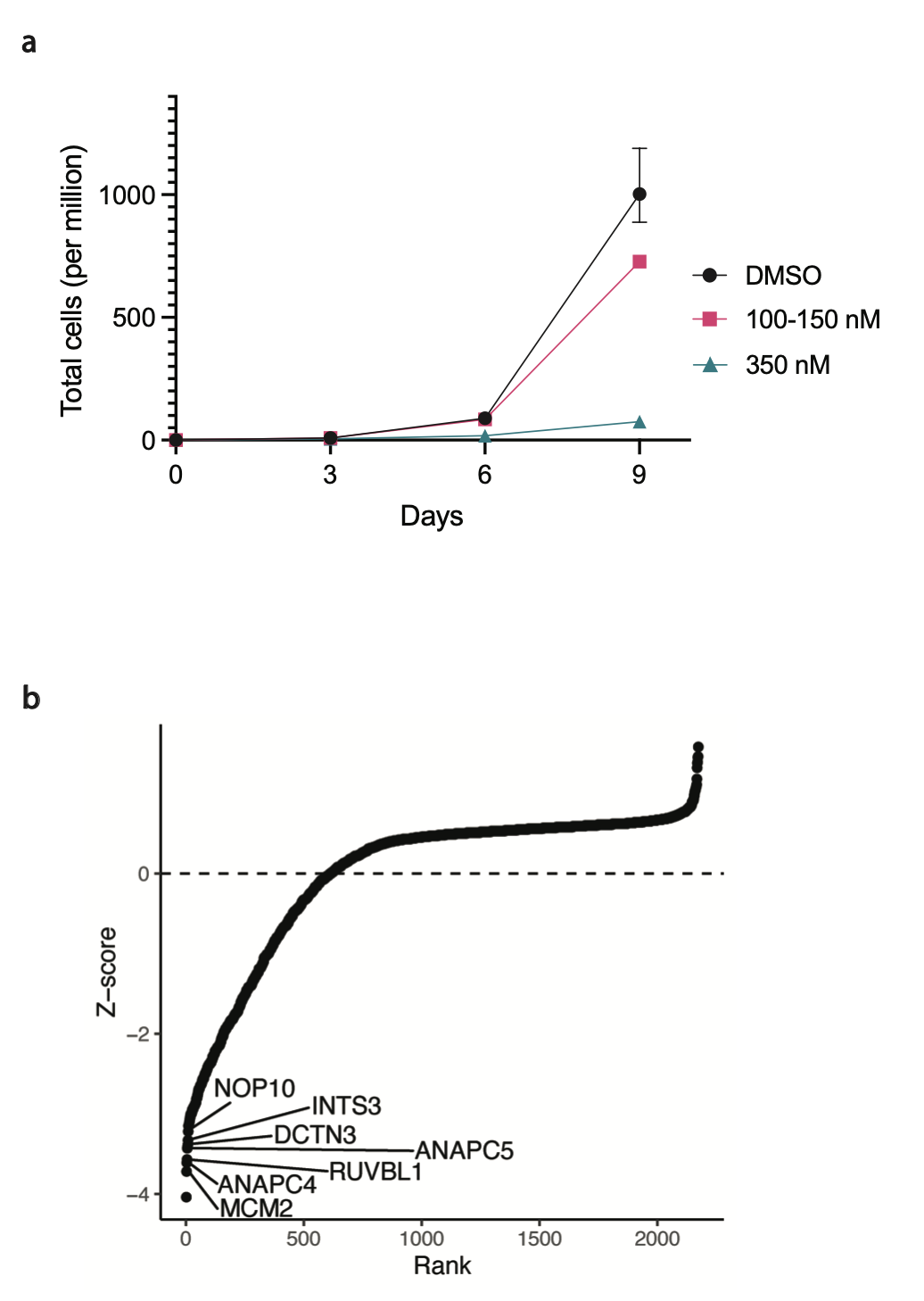
**Supplementary Material**

**Supplementary Fig.1: CRISPRi screen dosing and essentialome. a** Graph showing the growth over time of RPE TP53^-/-^ dCas9-KRAB cells treated with either DMSO or AZD1775. The 100-150 nM AZD1775 arm was treated with 100 nM for days 0-6 and treated with 150 nM on days 6-9 (biological n=2). Graphs are depicted with means ± SD. **b** Graph showing the NormZ score of sgRNAs on day 18 (DMSO treated arm) post-transduction compared to day 3 post-transduction. A handful of essential genes have been labelled to show a decrease of sgRNA representation on day 18 post-transduction.

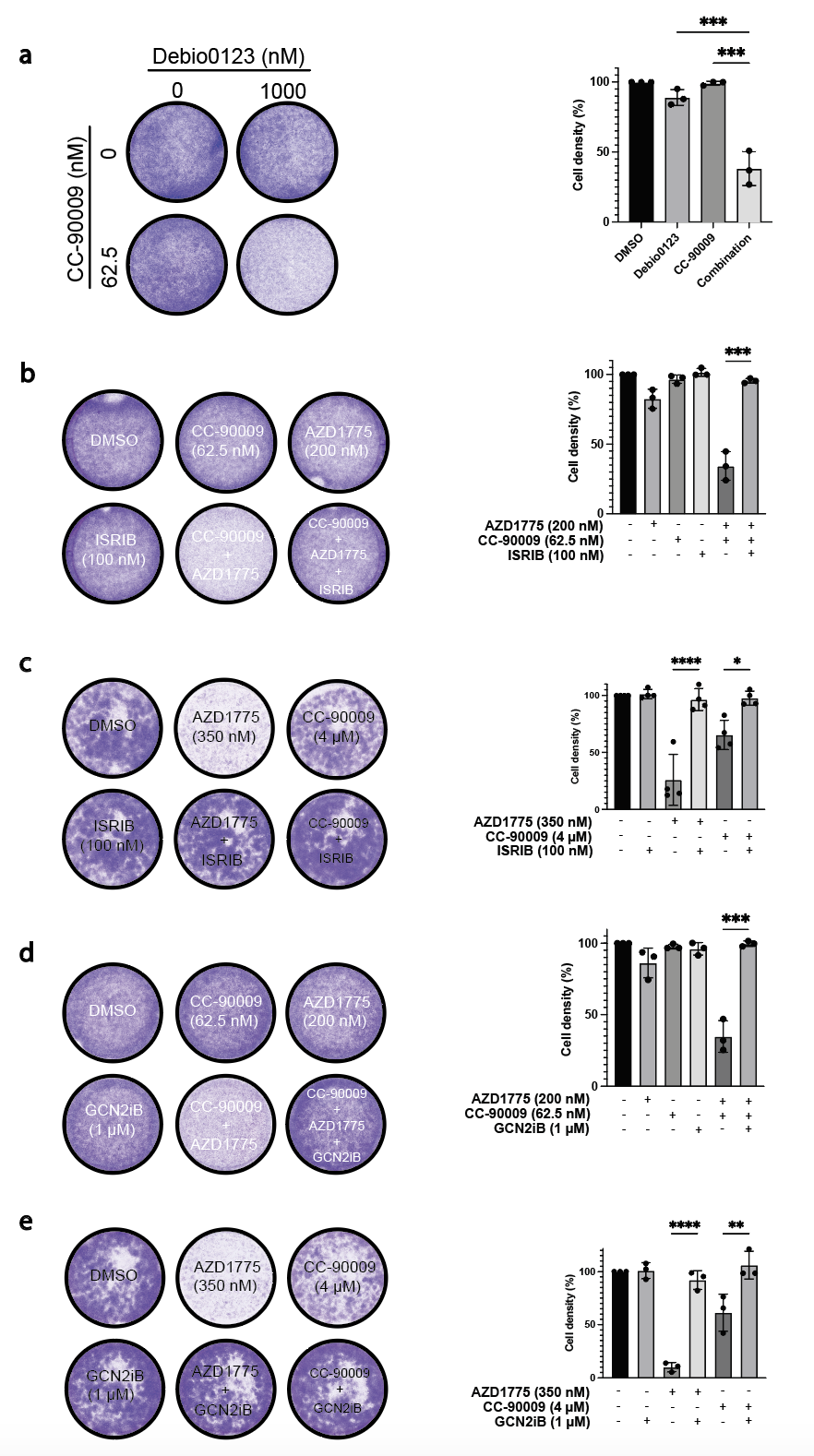

**Supplementary Fig.2: 6 well plate crystal violet assays on the RPE TP53^-/-^ cell line.** Representative images and graphs showing the RPE TP53^-/-^ cell line treated with DMSO, WEE1 inhibitors, ISRIB, GCN2iB alone and in combination. Following their respective timepoints, 6 well plates were washed and stained with crystal violet (biological n=3 with exception of c which is n=4). Graphs are depicted with means ± SD, points represent each biological replicate. Statistical analyses were performed using a one-way ANOVA test, * *p* < 0.05, ** *p* < 0.01, *** *p* < 0.001, **** *p* < 0.0001. **a, b, d** Cells were treated for 72 hours. **c, e** Cells were treated for 6 days. After 72 hours, fresh media and drug was applied.

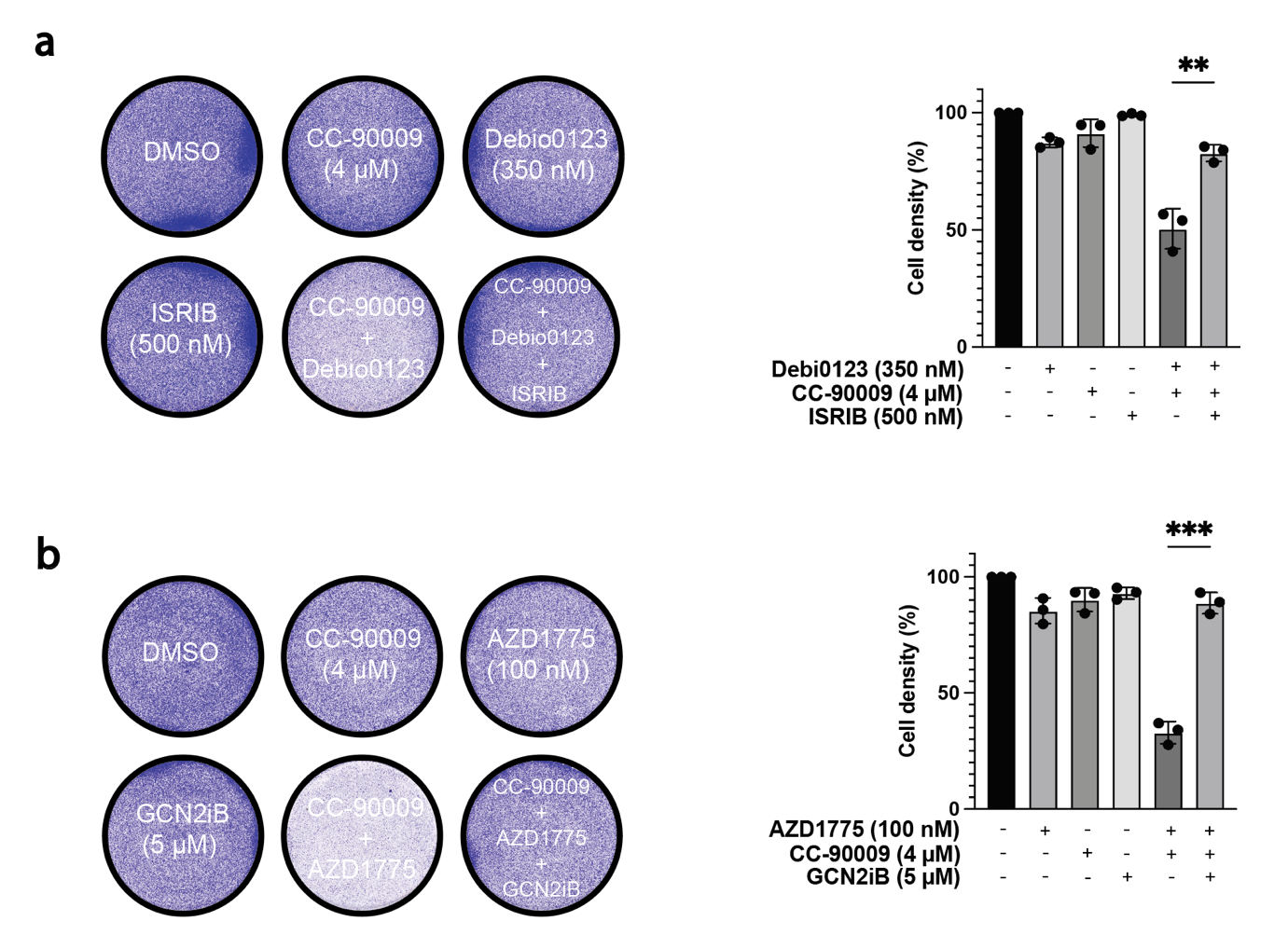

**Supplementary Fig.3: 6 well plate crystal violet assays on the HAP1 cell line. a, b** Representative images and graphs showing the HAP1 cell line treated with DMSO, WEE1 inhibitors, CC-90009, ISRIB, GCN2iB alone and in combination for 72 hours. Following their respective timepoints, 6 well plates were washed and stained with crystal violet (biological n=3). Graphs are depicted with means ± SD, points represent each biological replicate. Statistical analyses were performed using a one-way ANOVA test, ** *p* < 0.01, *** *p* < 0.001.

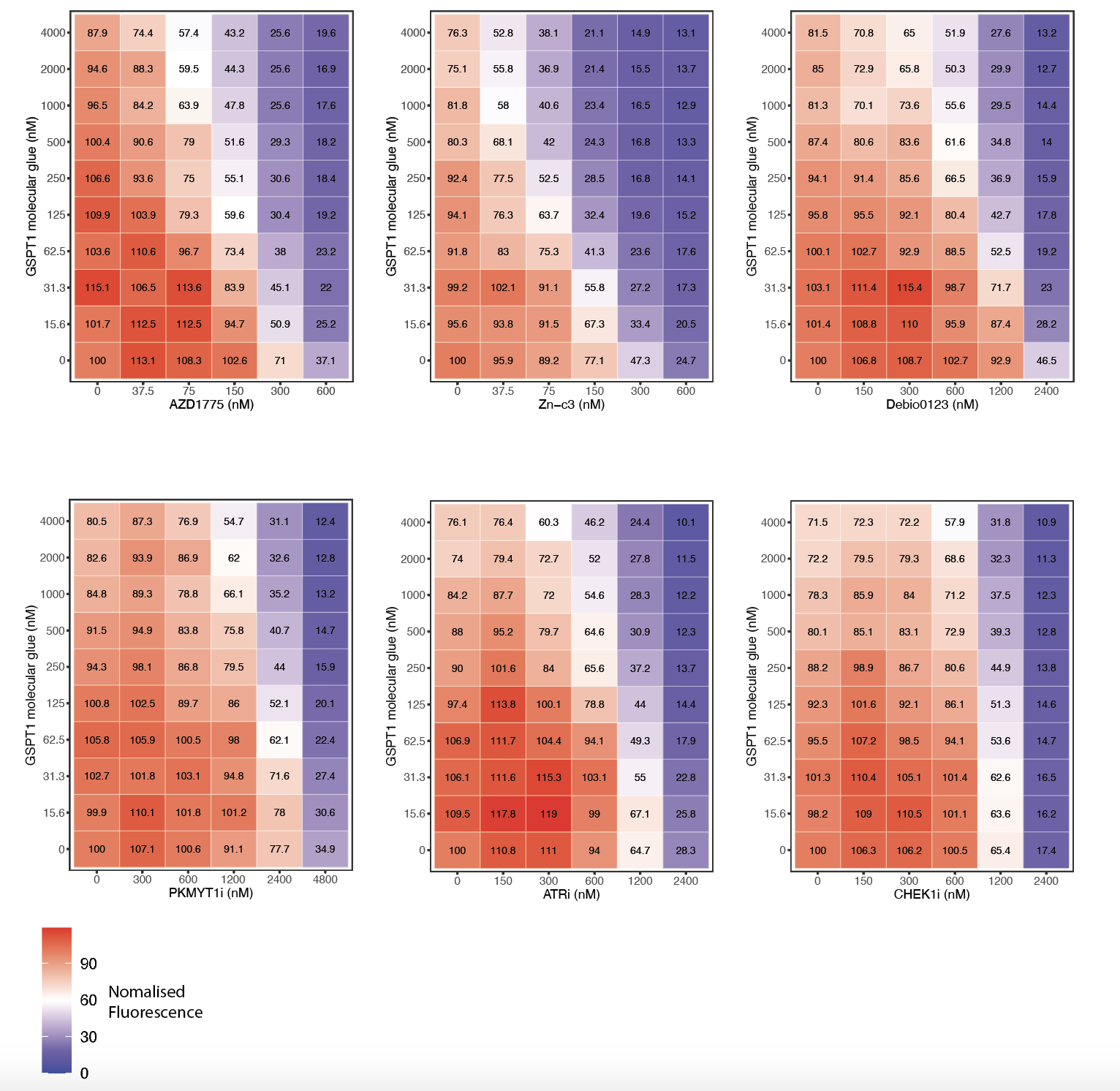

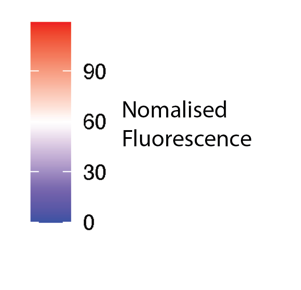

**Supplementary Fig.4: Heatmaps of CC-90009-DNA damage response inhibitor combinations.** Heatmaps showing resazurin cell viability assays in a 96 well plate format in the RPE TP53^-/-^ cell line. CC-90009 was treated in combination with WEE1 inhibitors (AZD1775, Debio0123 and Zn-c3). PKMYT1i (RP-6306), ATRi (AZD6738) or CHEK1i (LY2603618) for 72 hours (biological n=4).

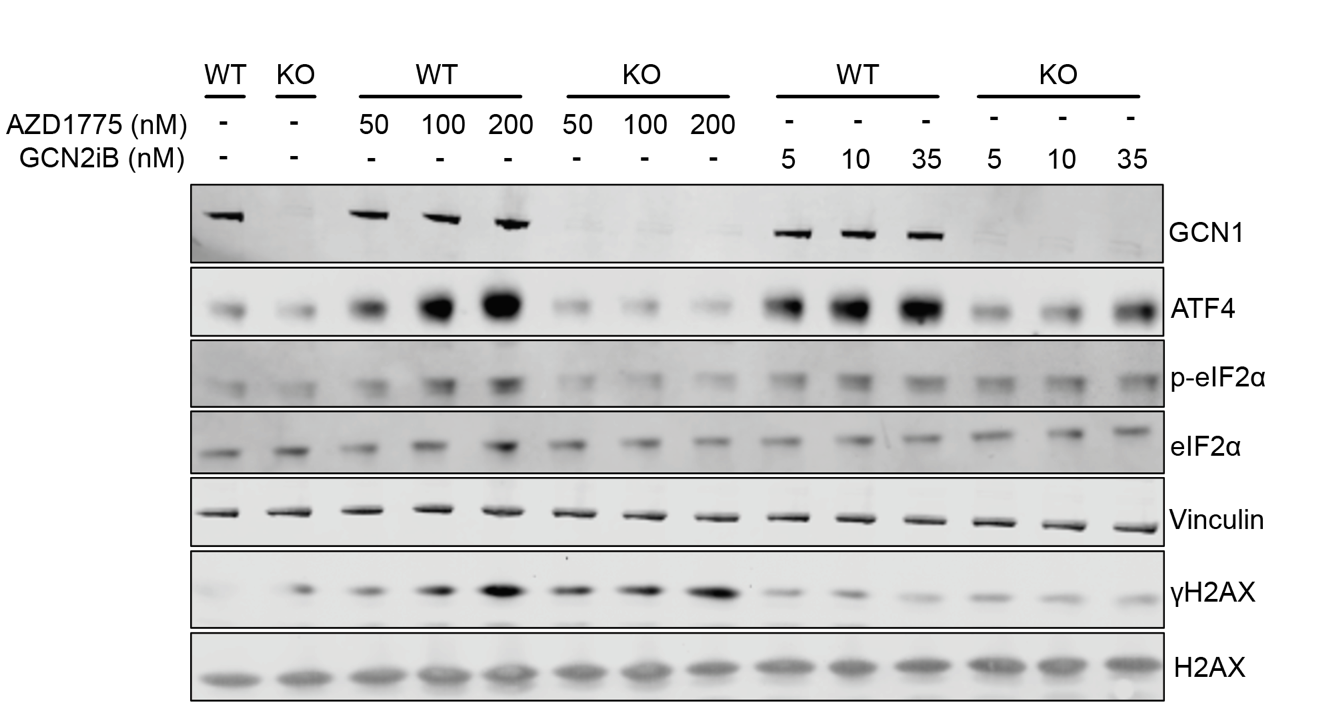

**a**

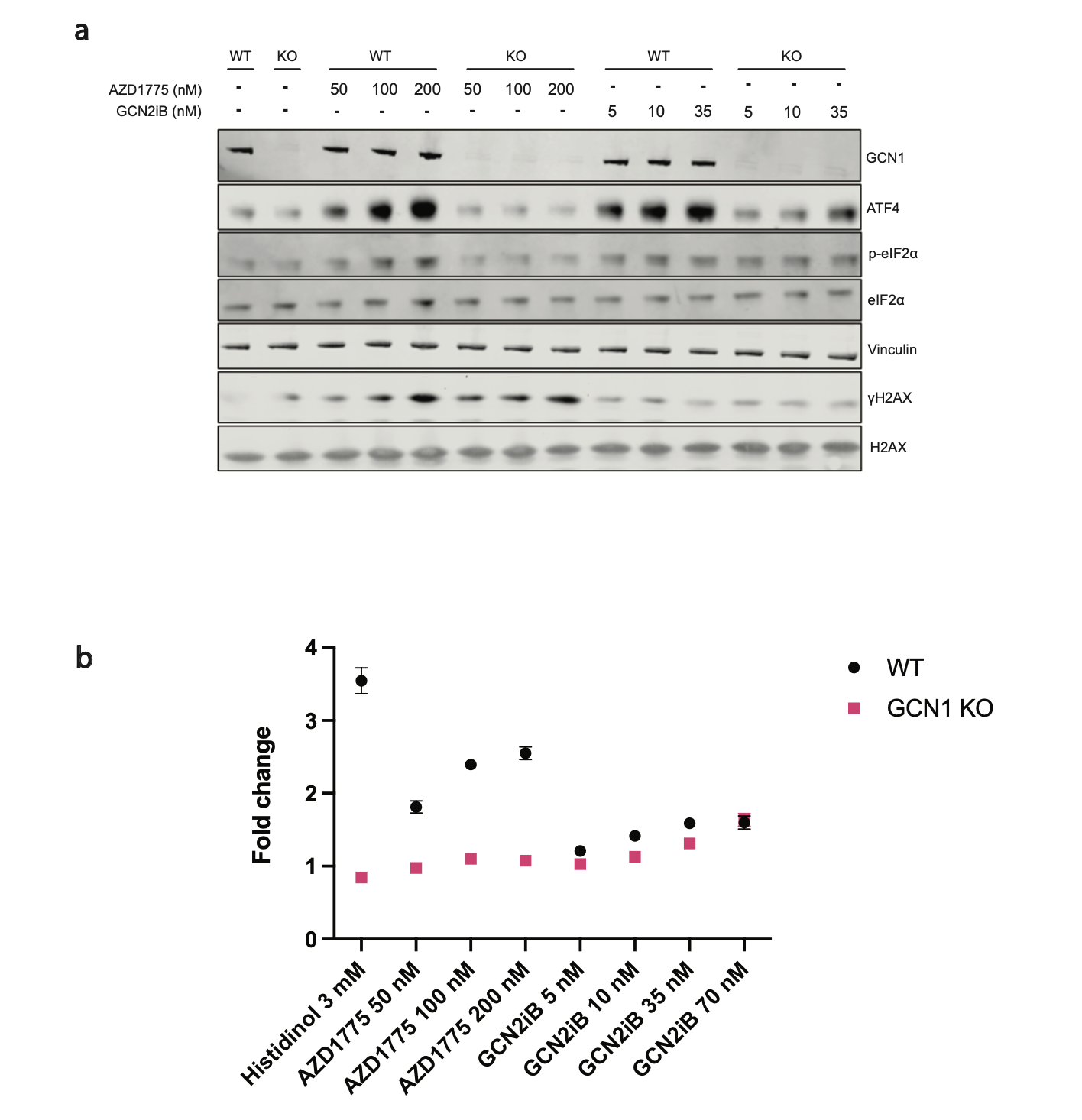

**b**

**Supplementary Fig.5: Reduced ISR activation from AZD1775 in the HEK293 *GCN1* knock-out cell line compared to wildtype. a** Western blot showing 6 hour treatments of HEK293 wild type and *GCN1* knockout cell lines. **b** A graph showing the fold change of ATF4 reporter^1^ signal that was transfected into HEK293 wildtype and GCN1 knockout cell lines. Transfected cells were treated for 6 hours with either histidinol, AZD1775 or GCN2iB (biological n=3, except for 70 nM GCN2iB which is biological n=2). Graphs are depicted with means ± SEM.

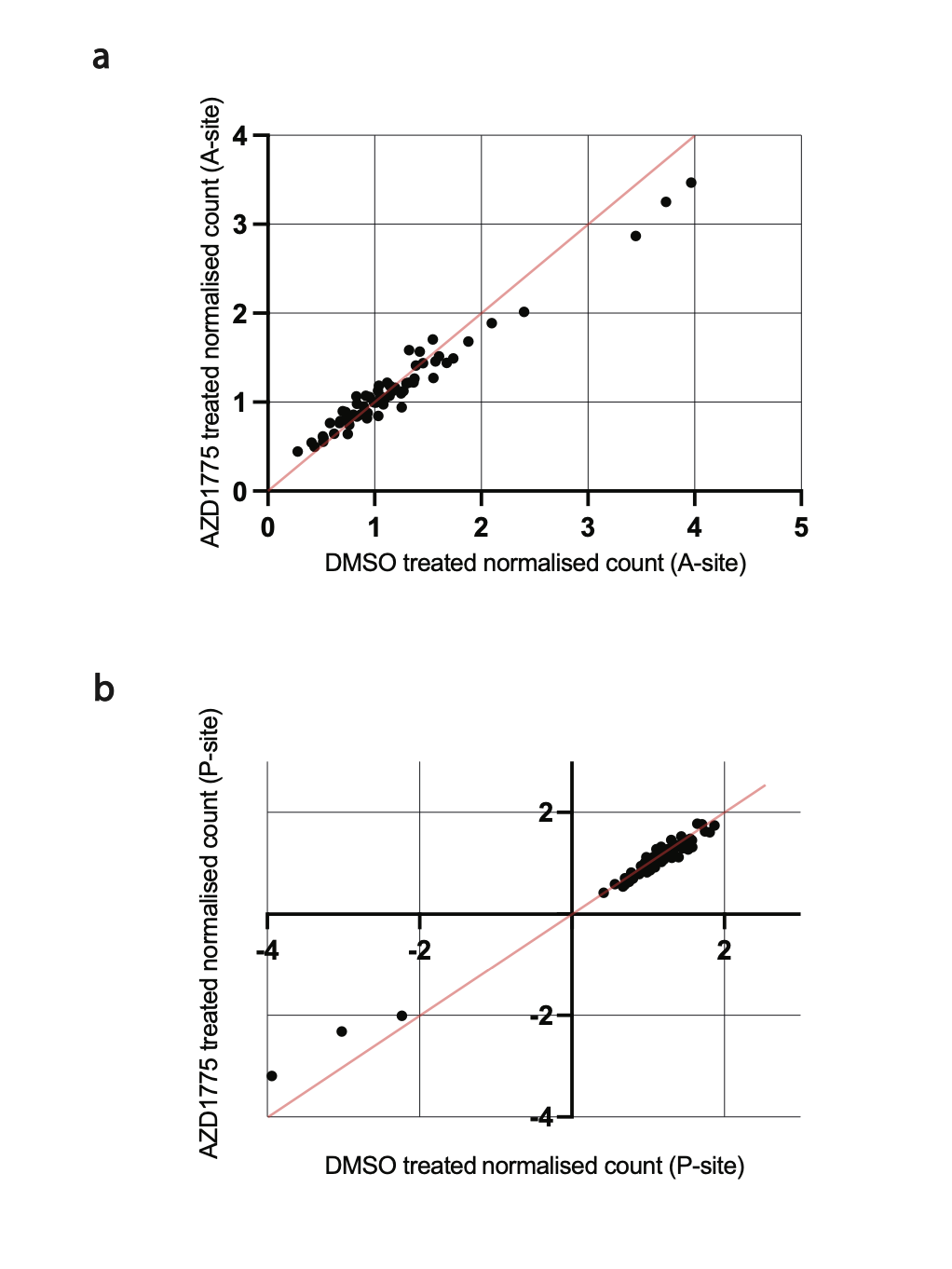

**a**

**b**

**Supplementary Fig.6: A comparison of the ribosome occupancy of AZD1775 treatment vs DMSO. a, b** Graphs showing the codon occupancy in the A site and P site of the ribosome respectively. DMSO or 650 nM AZD1775 were treated on the RPE TP53^-/-^ cell line for 10 hours. Each point represents a particular codon (biological n=3).

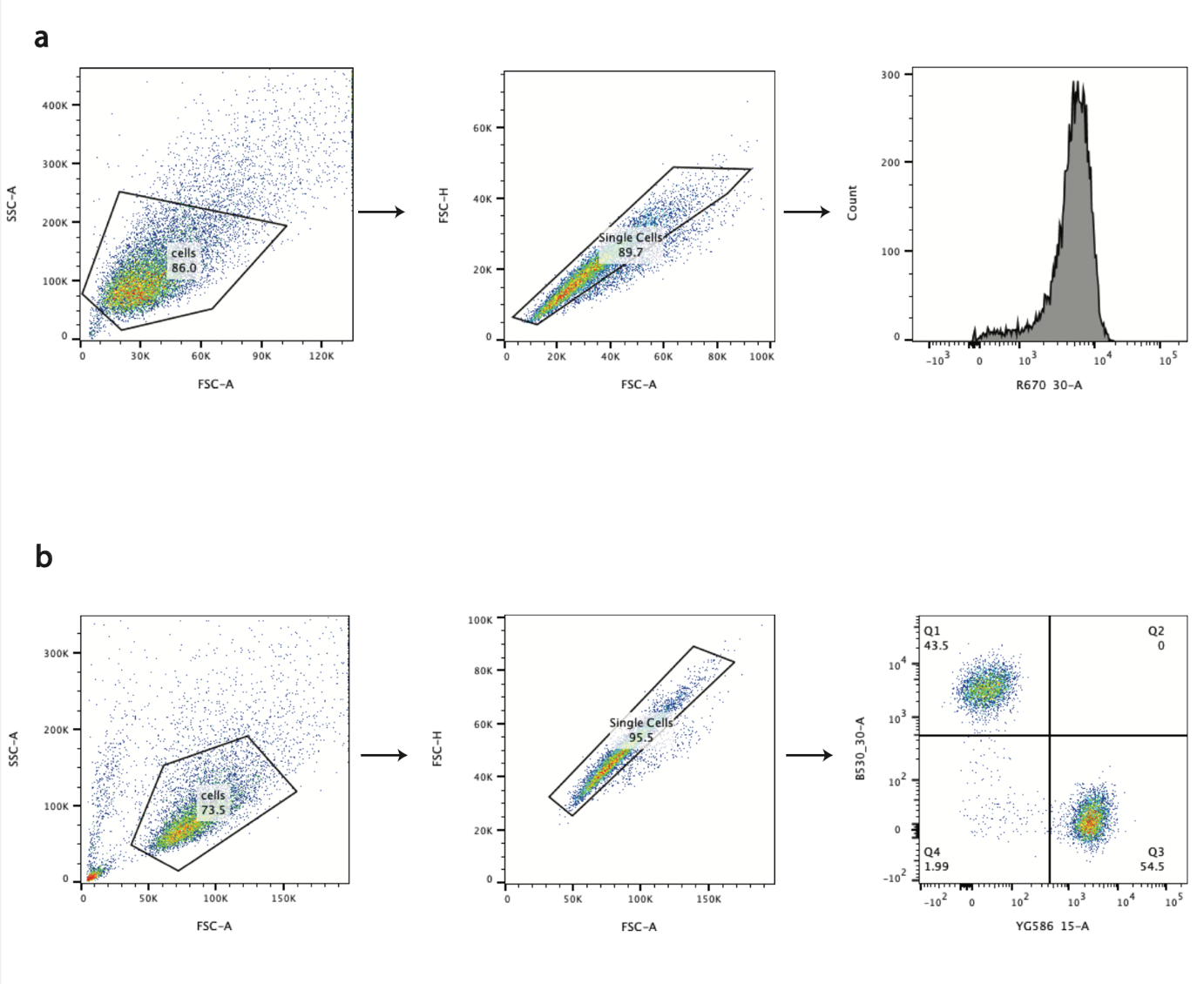
**a**

**b**
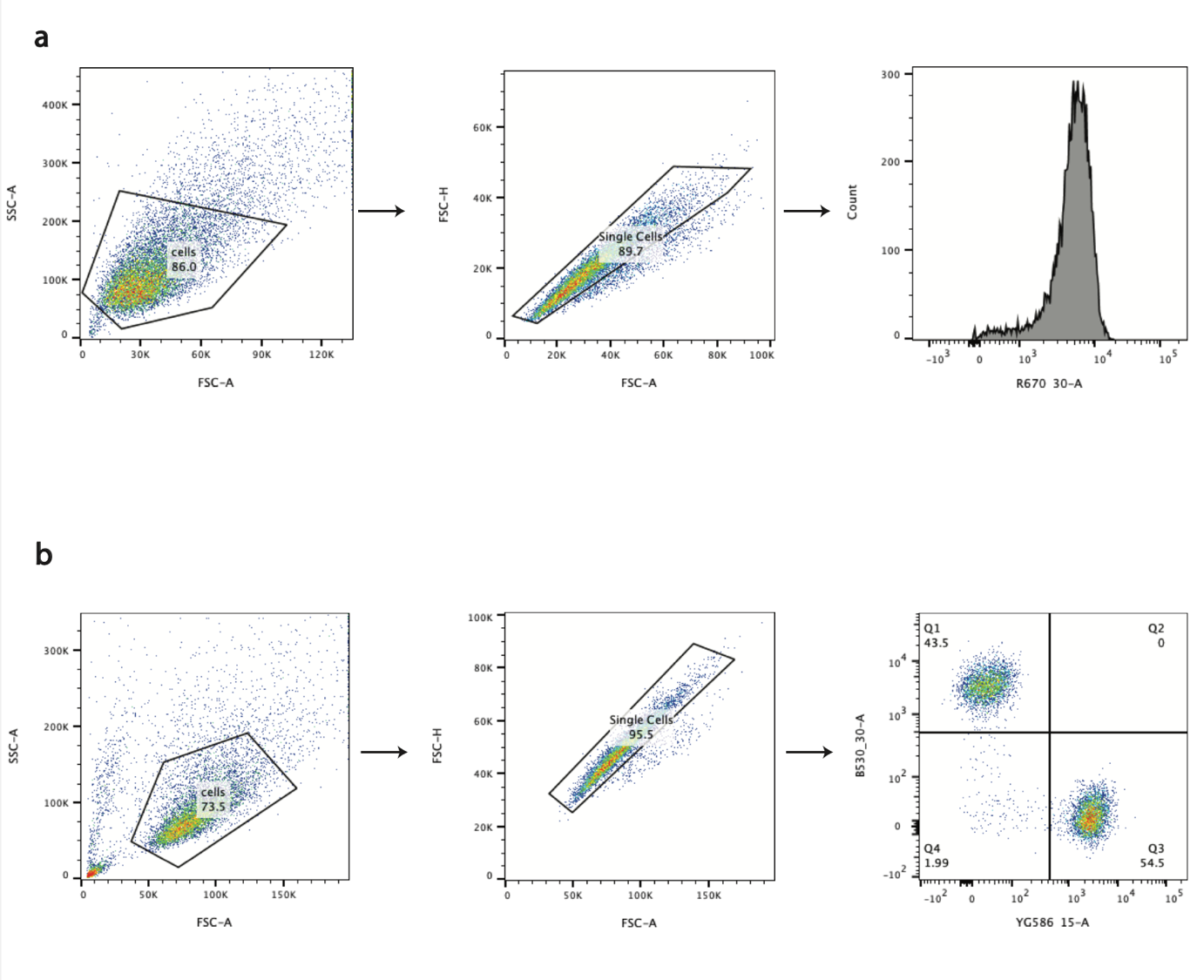

**Supplementary Fig.7: Flow cytometry gating strategies. a, b** Flow cytometry gating for the AHA click reaction in the RPE TP53^-/-^ cell line and for the CRISPRi-based two colour growth competition assays in the RPE TP53^-/-^ dCas9-KRAB cell line to quantify the GFP: mCherry ratios respectively.

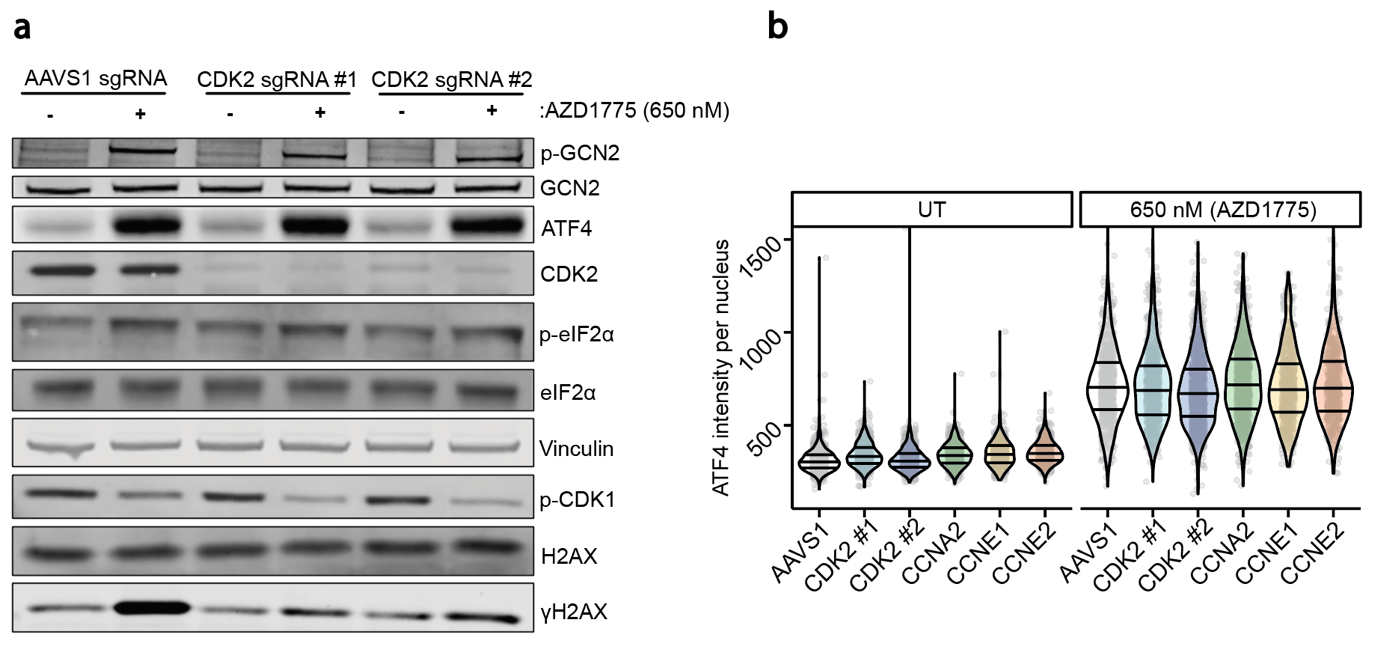

**Supplementary Fig.8: The depletion of CDK2, cyclin A2, cyclin E1 or cyclin E2 does not impact WEE1i induced ISR. a** Western blot showing the RPE TP53^-/-^ dCas9-KRAB cell line expressing an sgRNA that targets CDK2 or the AAVS1 locus treated with DMSO or 650 nM AZD1775 for 24 hours. **b** Immunofluorescence probing for nuclear ATF4 in the RPE TP53^-/-^ dCas9-KRAB cell line expressing a sgRNA that targets CDK2 (sgRNA #1 and #2), cyclin A2, cyclin E1 or cyclin E2 or the AAVS1 locus. Cells were treated with DMSO or 650 nM AZD1775 for 24 hours (biological n=3). Violin plots show median and quartile ranges.

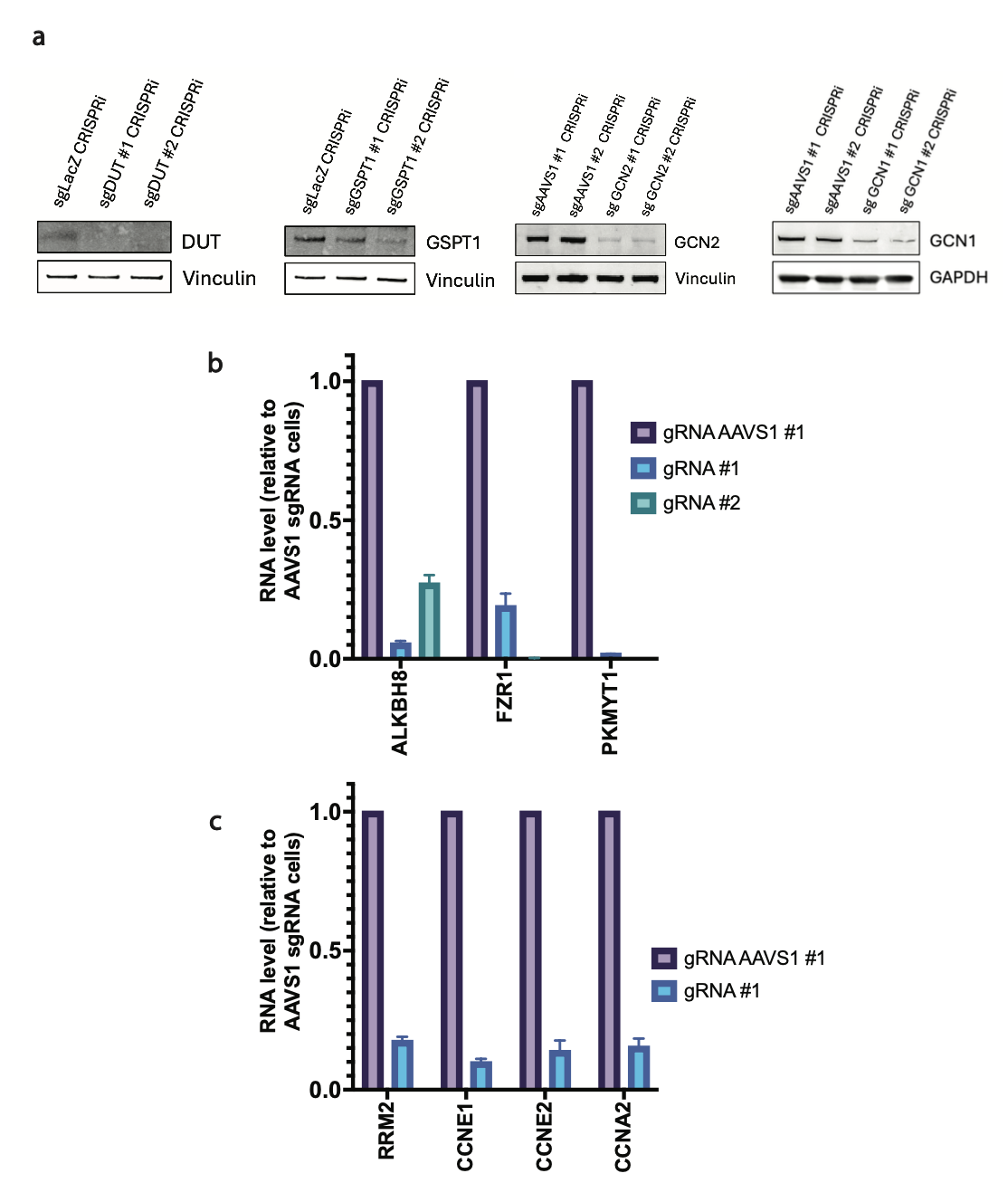
**Supplementary Fig.9: Validations for CRISPRi mediated knockdowns in the RPE TP53^-/-^ dCas9-KRAB cell line. a** Western blots showing RPE TP53^-/-^ dCas9-KRAB cell lines expressing sgRNAs that target DUT, GSPT1, GCN2 and GCN1 compared to their respective controls that are either non-targeting (LacZ) or targeting the AAVS1 locus. **b, c** RT-qPCRs of RPE TP53^-/-^ dCas9-KRAB cell line expressing sgRNAs that target ALKBH8, RRM2, FZR1, PKMYT1, CCNE1, CCNE2 and CCNA2. RNA level was relative to the RPE TP53^-/-^ dCas9-KRAB cell line expressing an sgRNA that target the AAVS1 locus. Measuring the GAPDH RNA abundance for each cell line was used as a control (technical n=3). Graphs are depicted with means ± SD.

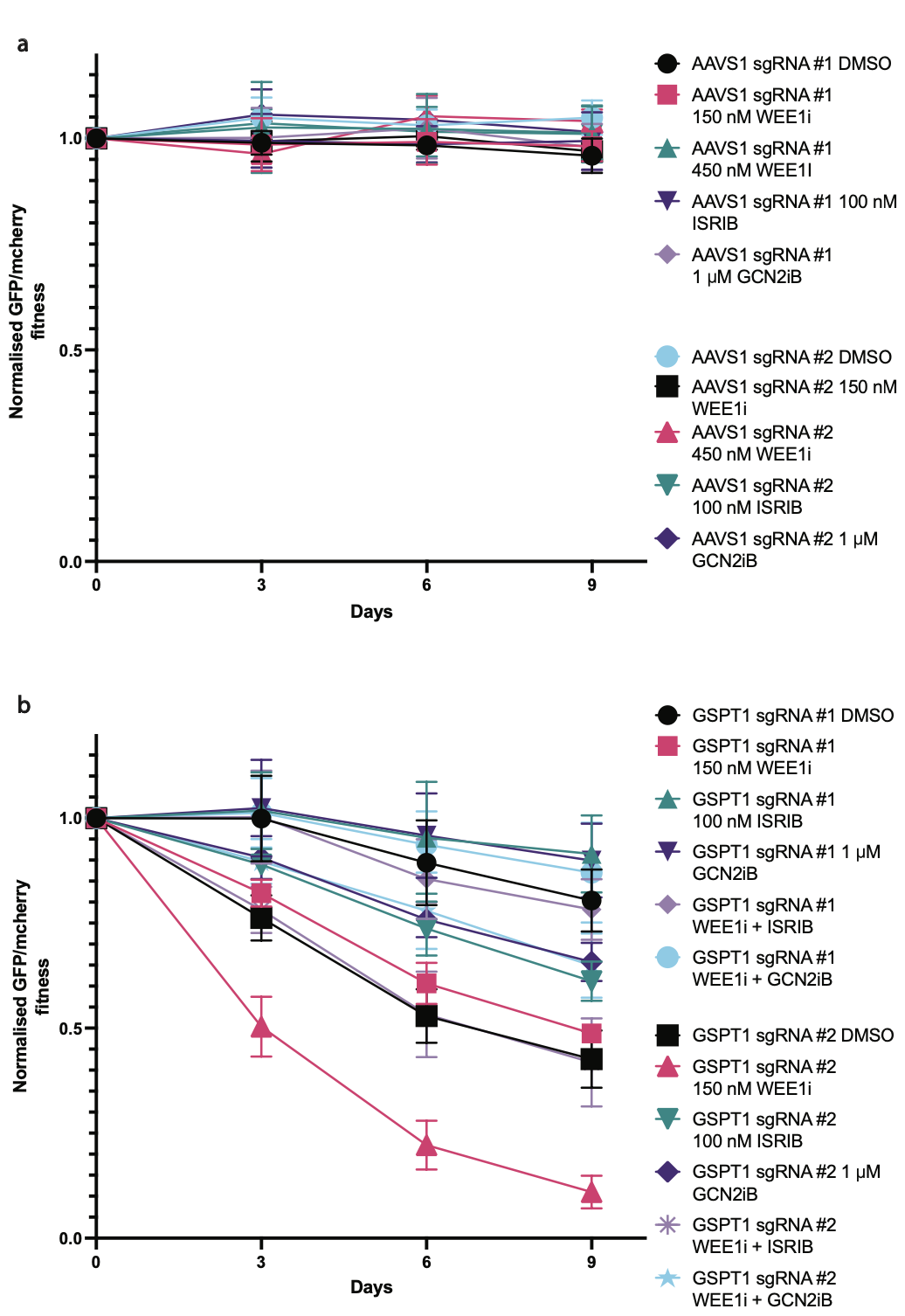

**Supplementary Fig.10: Normalised GFP/mCherry fitness graphs for sgAAVS1-GFP vs sgLacZ-mCherry and sgGSPT1-GFP vs sgLacZ-mCherry. a, b** RPE TP53^-/-^ dCas9-KRAB cells expressing sgRNA of interest and GFP were mixed 50/50 with RPE TP53^-/-^ dCas9-KRAB cells expressing sgLacZ-mCherry on day 0. Cells were passaged in a 12 well plate format every 3 days and fresh drug was added. The GFP and mCherry abundance was measured every 3 days for 9 days. Values above 1 indicate the cell population expressing GFP had an increased relative cell fitness compared to cells expressing mCherry; whereas values below 1 indicate the cell population expressing GFP had a reduction in relative cell fitness compared to cells expressing mCherry (biological n=3). Graphs are depicted with means ± SD.

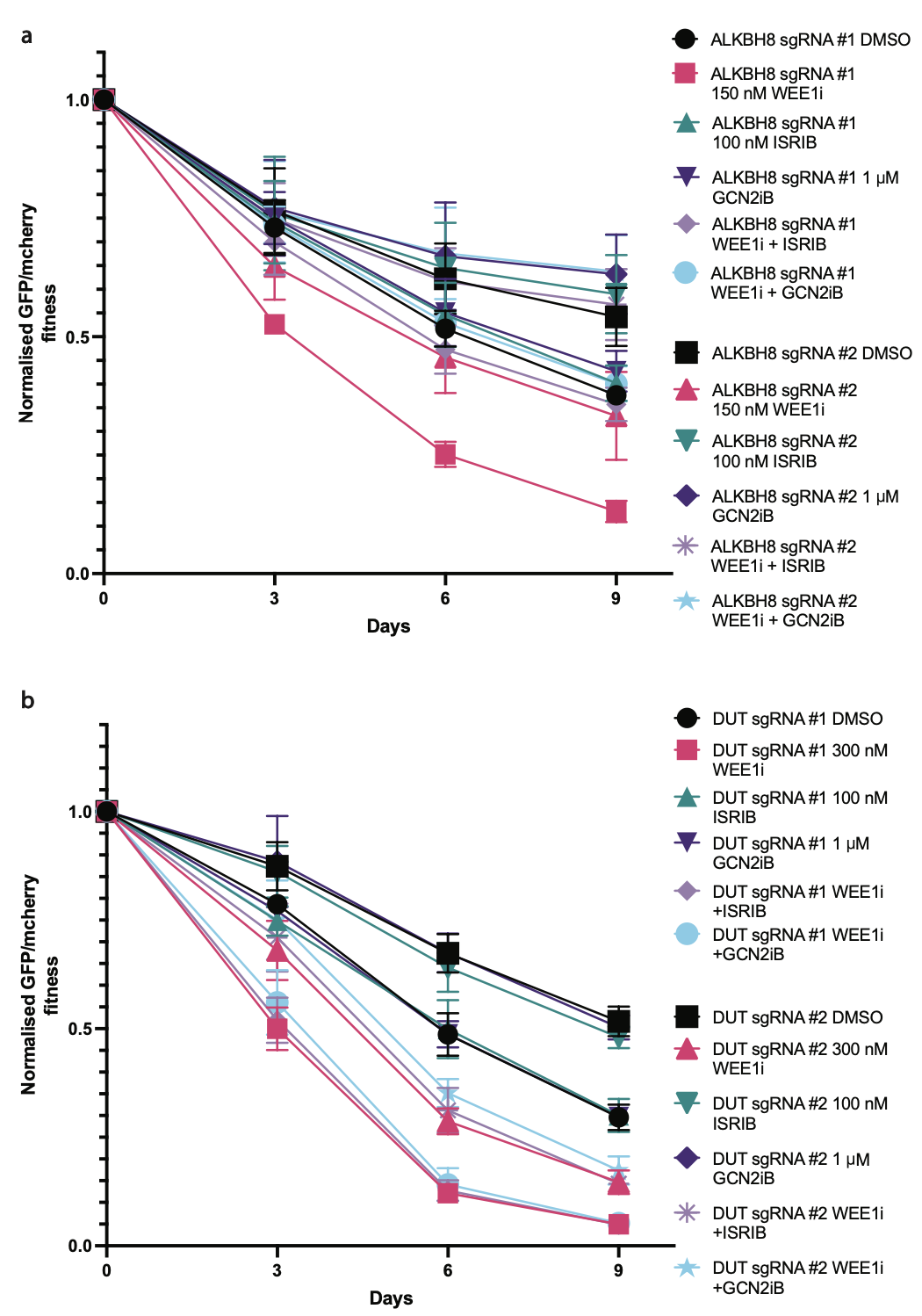

**Supplementary Fig.11: Normalised GFP/mCherry fitness graphs for sgALKBH8-GFP vs sgLacZ-mCherry and sgDUT-GFP vs sgLacZ-mCherry. a, b** RPE TP53^-/-^ dCas9-KRAB cells expressing sgRNA of interest and GFP were mixed 50/50 with RPE TP53^-/-^ dCas9-KRAB cells expressing sgLacZ-mCherry on day 0. Cells were passaged in 12 well plate format every 3 days and fresh drug was added. The GFP and mCherry abundance was measured every 3 days for 9 days. Values above 1 indicate the cell population expressing GFP had an increased relative cell fitness compared to cells expressing mCherry; whereas values below 1 indicate the cell population expressing GFP had a reduction in relative cell fitness compared to cells expressing mCherry (biological n=3). Graphs are depicted with means ± SD.

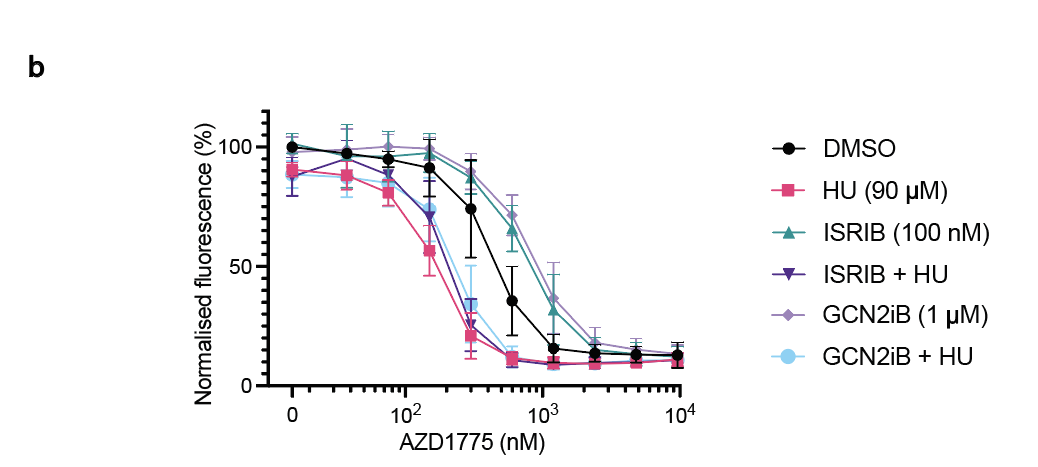

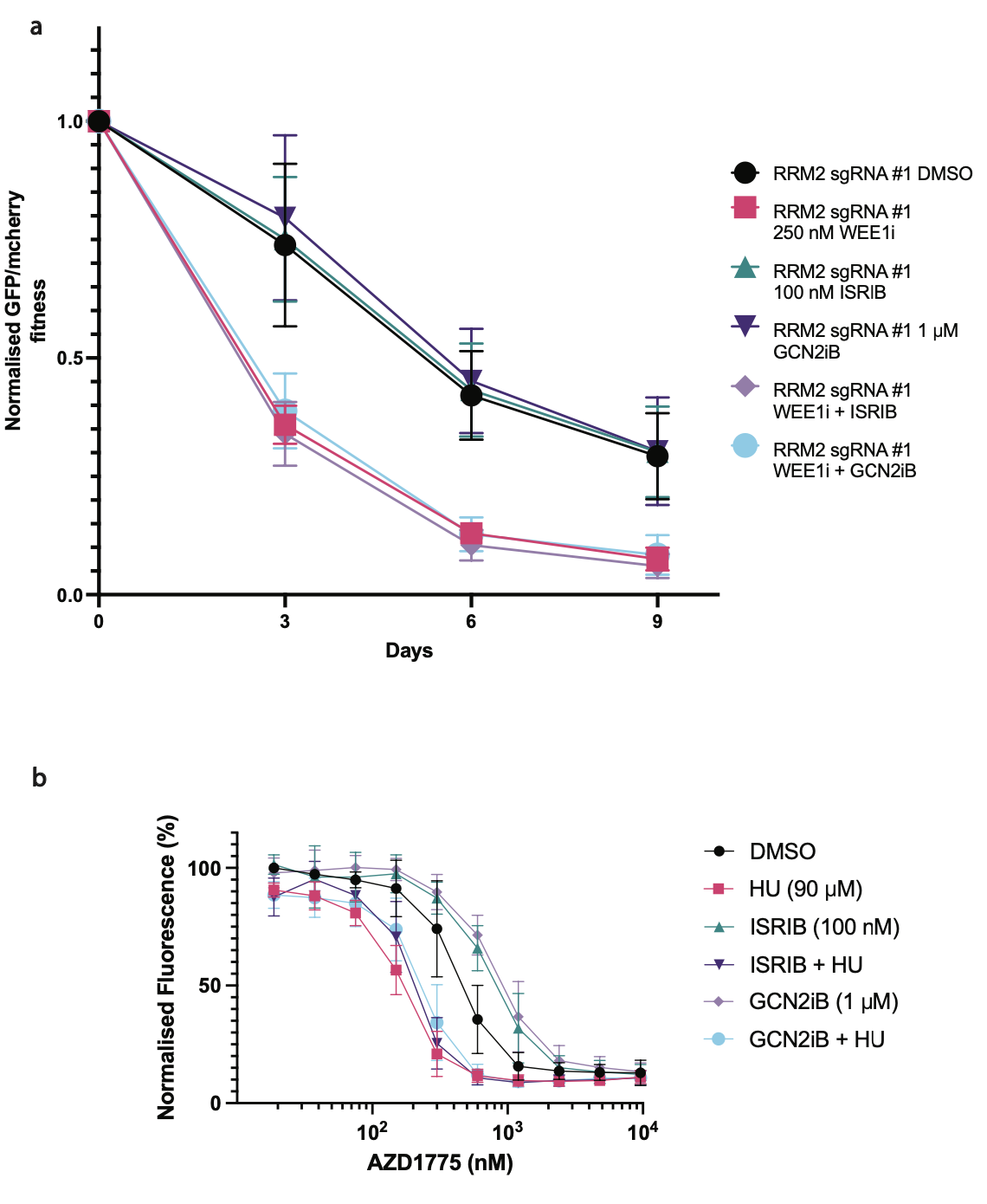

**Supplementary Fig.12: Normalised GFP/mCherry fitness graph for sgRRM2-GFP vs sgLacZ-mCherry and cell viability assay of hydroxyurea in combination with AZD1775. a** RPE TP53^-/-^ dCas9-KRAB cells expressing sgRRM2 and GFP were mixed 50/50 with RPE TP53^-/-^ dCas9-KRAB cells expressing sgLacZ-mCherry on day 0. Cells were passaged in 12 well plate format every 3 days and fresh drug was added. The GFP and mCherry abundance was measured every 3 days for 9 days. Values above 1 indicate the cell population expressing GFP had an increased relative cell fitness compared to cells expressing mCherry; whereas values below 1 indicate the cell population expressing GFP had a reduction in relative cell fitness compared to cells expressing mCherry (biological n=3). Graphs are depicted with means ± SD. **b** Resazurin cell viability assay with varying concentrations of AZD1775 with and without DMSO, 90 μM hydroxyurea, 100 nM ISRIB, 100 nM ISRIB + 90 μM hydroxyurea, 1 μM GCN1iB and 1 μM GCN1iB + 90 μM hydroxyurea treated on the RPE TP53^-/-^ cell line for 72 hours (biological n=4). Graphs are depicted with means ± SD.

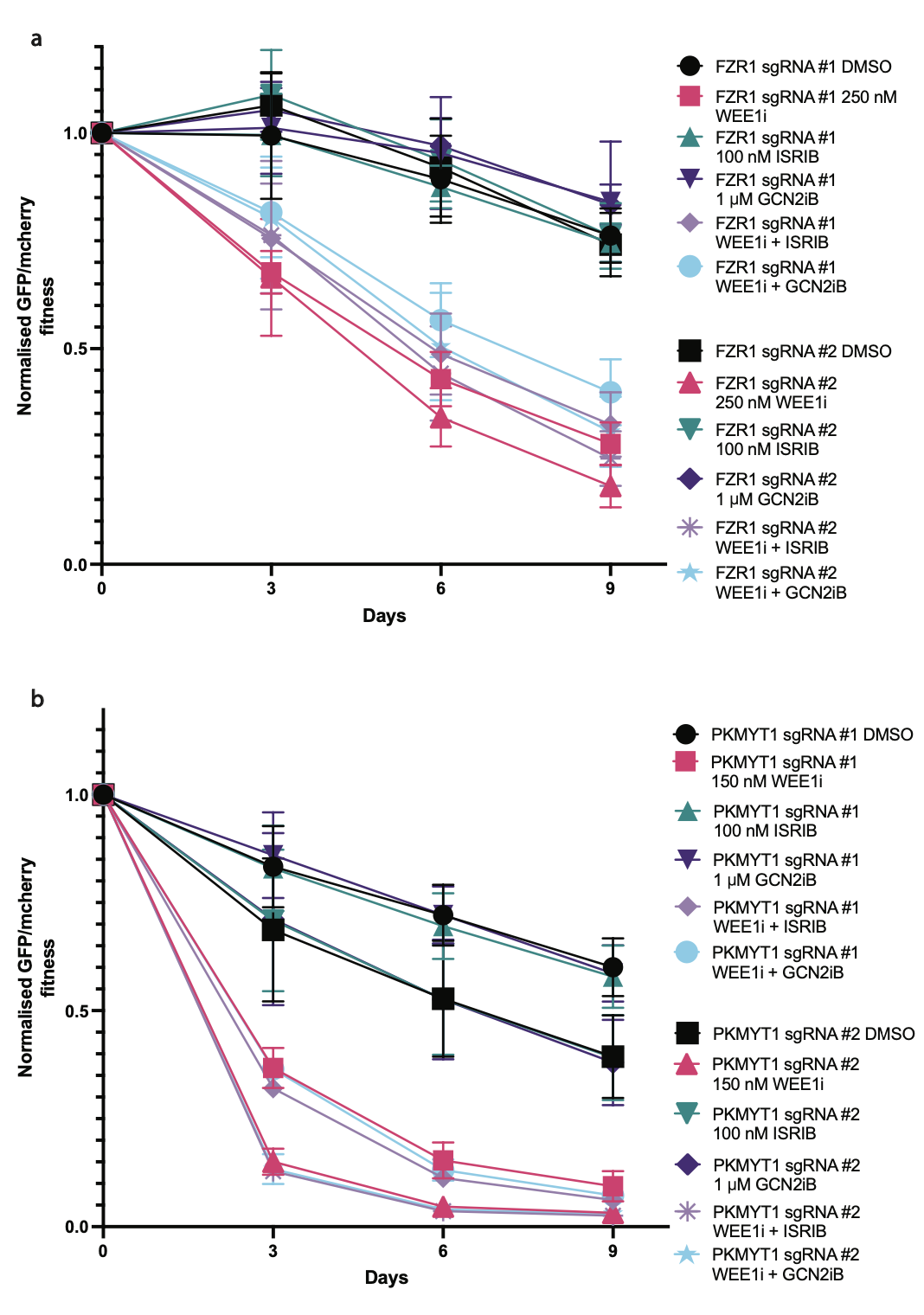

**Supplementary Fig.13: Normalised GFP/mCherry fitness graphs for sgFZR1-GFP vs sgLacZ-mCherry and sgPKMYT1-GFP vs sgLacZ-mCherry. a, b** RPE TP53^-/-^ dCas9-KRAB cells expressing sgRNA of interest and GFP were mixed 50/50 with RPE TP53^-/-^ dCas9-KRAB cells expressing sgLacZ-mCherry on day 0. Cells were passaged in 12 well plate format every 3 days and fresh drug was added. The GFP and mCherry abundance was measured every 3 days for 9 days. Values above 1 indicate the cell population expressing GFP had an increased relative cell fitness compared to cells expressing mCherry; whereas values below 1 indicate the cell population expressing GFP had a reduction in relative cell fitness compared to cells expressing mCherry (biological n=3). Graphs are depicted with means ± SD.

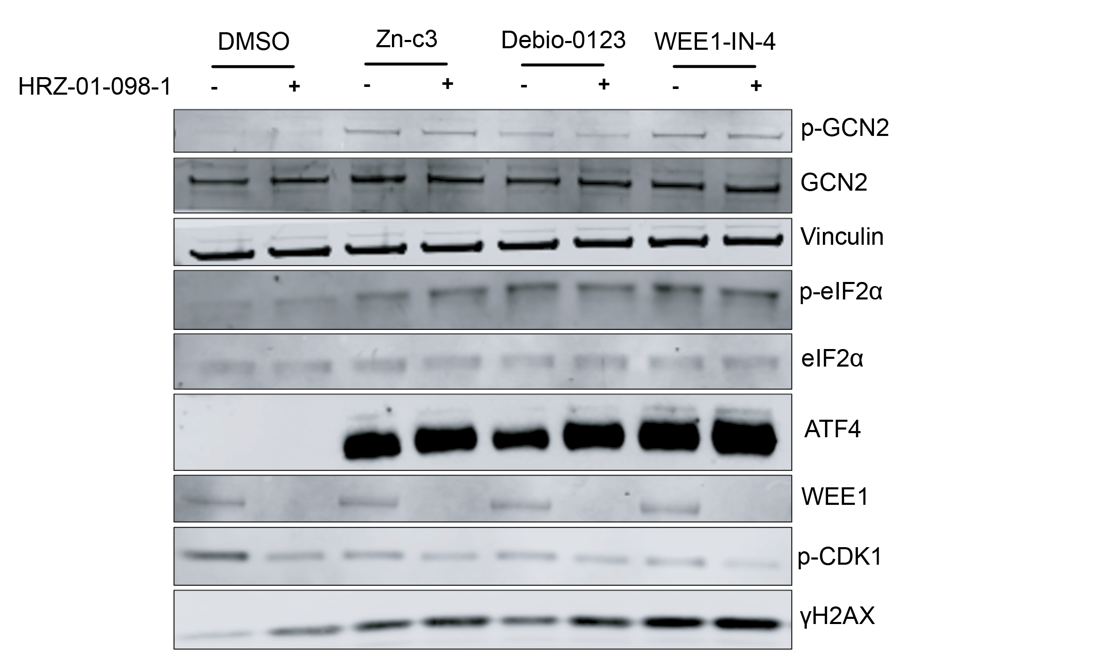

**a**

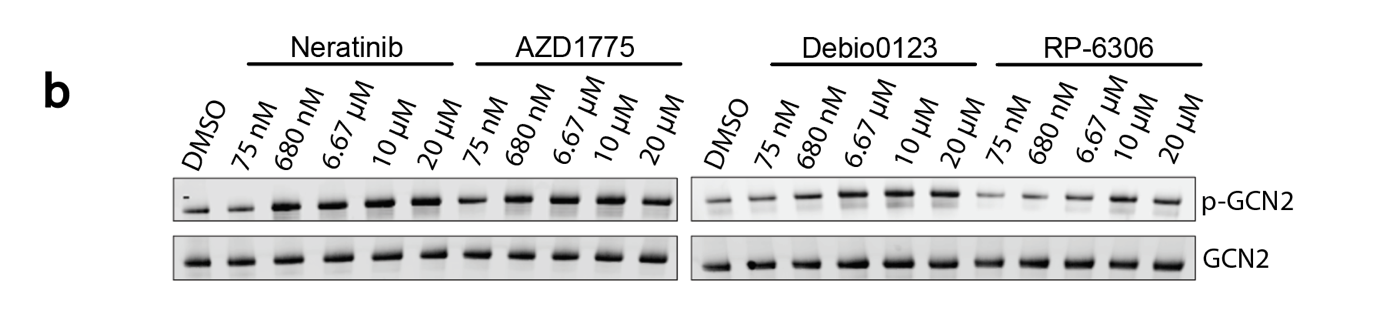

**b**

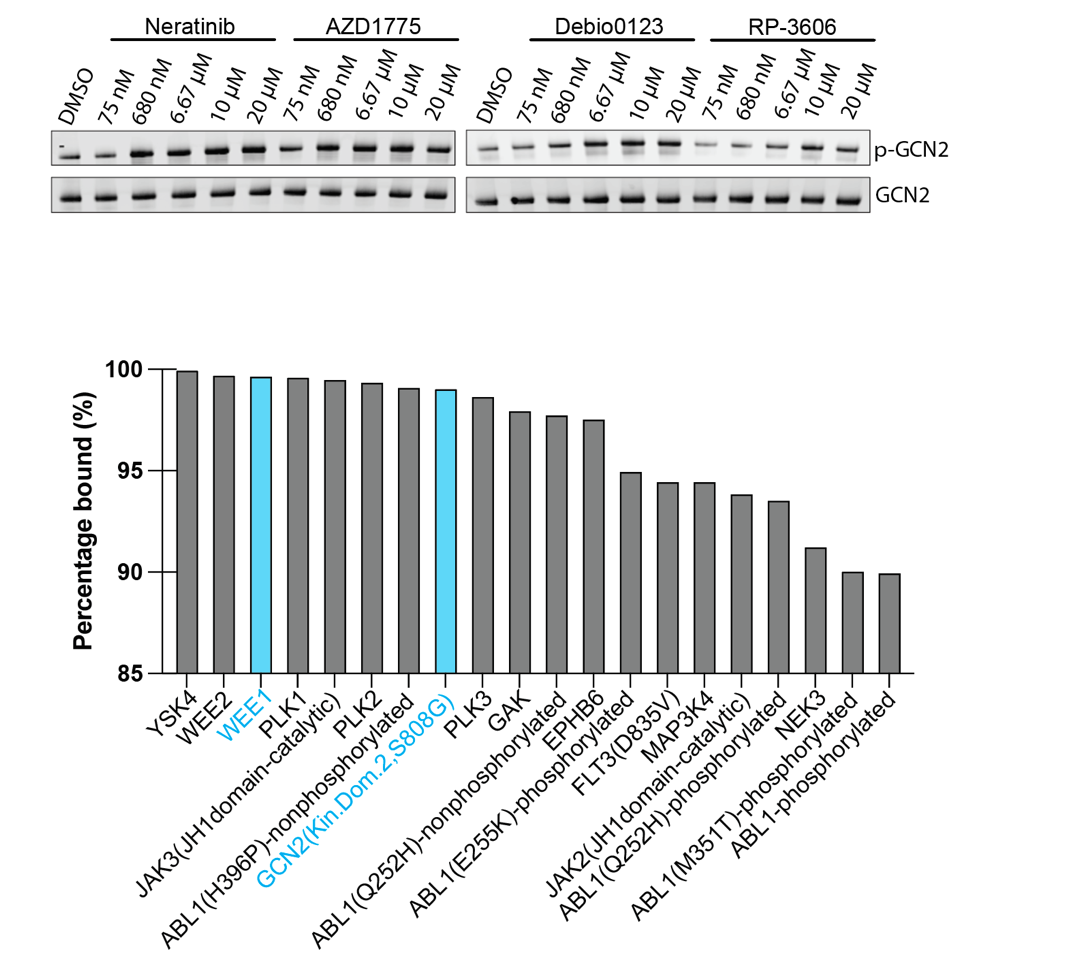

**c**

**Supplementary Fig.14: WEE1i induced ISR is independent of WEE1. a** Western blot of the RPE TP53^-/-^ cell line. Cells were pre-treated with DMSO or 1 μM HRZ-1-098-1 WEE1 molecular glue for 1 hour (7-hour treatment in total) followed by DMSO, 650 nM Zn-c3, 3 μM Debio0123 or 3 μM WEE1-IN-4 for 6 hours. **b** A western blot of an in vitro experiment probing the total and phosphorylated GCN2 in the presence of DMSO, Neratinib, WEE1i (AZD1775 and Debio0123) and PKMYT1i (RP-6306). **c** A bar chart showing a rank plot of the top 20 kinases and kinase domains that bound with the 0.5 μM AZD1775 compound from kinome profiling. A total of 403 wild-type and 65 mutant kinases were scanned. Data generated in previous publication^2^. WEE1 and the second domain of GCN2/eIF2AK4 are highlighted in blue.

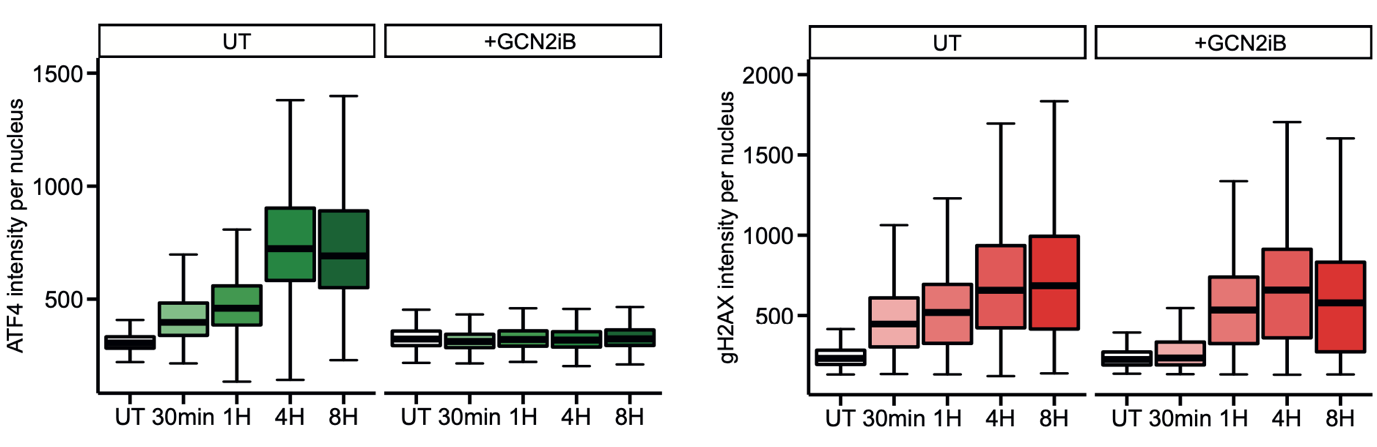

**Supplementary Fig.15: ISR induction but not γH2AX from AZD1775 treatment can be rescued by GCN2iB.** Immunofluorescence probing for nuclear ATF4 and γH2AX in the RPE TP53^-/-^ cell line. Cells were treated with 650 nM AZD1775 with and without 1 μM GCN2iB at different timepoints (biological n=3). Box plots show median and quartile ranges.

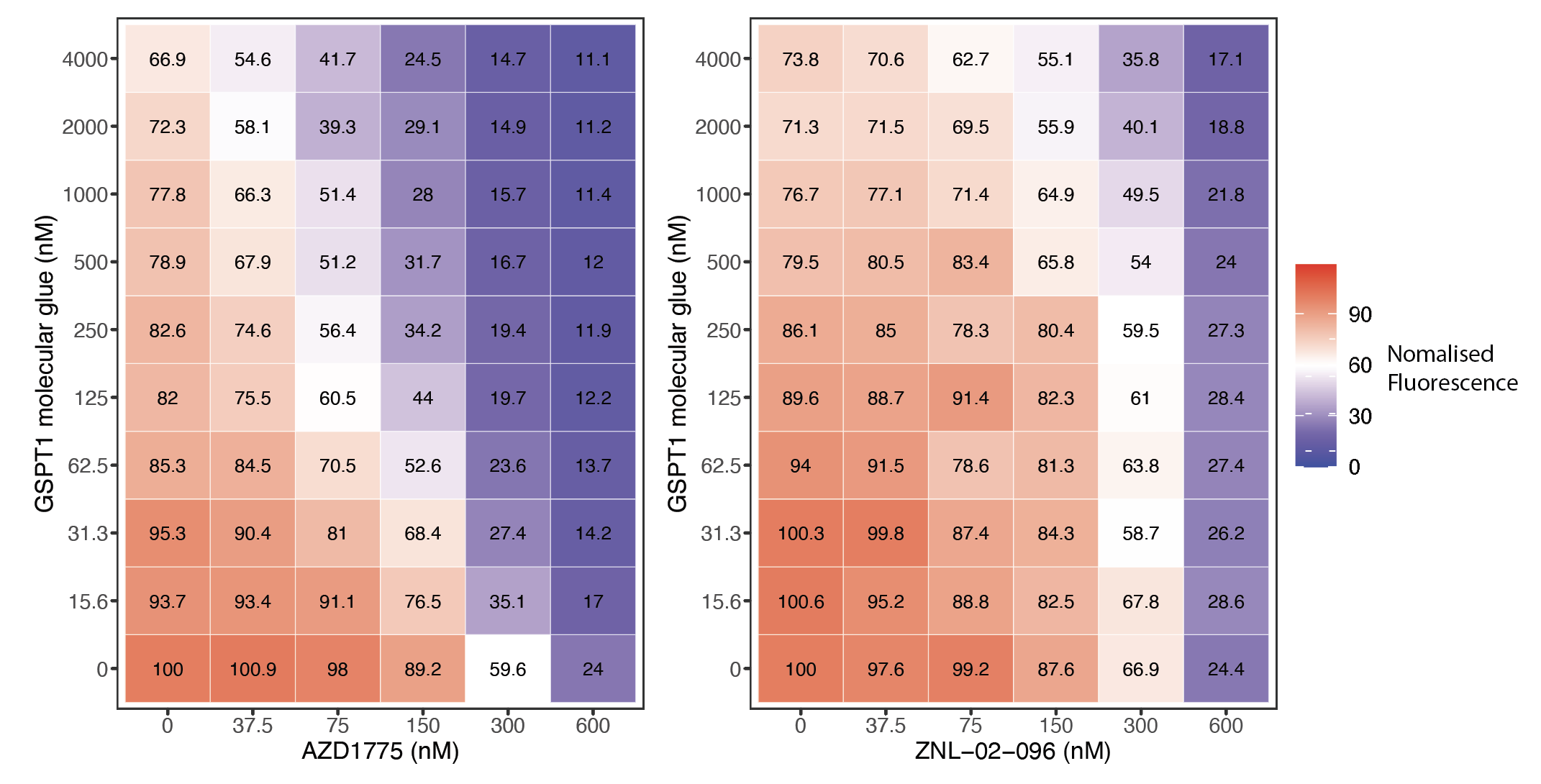
 **a**

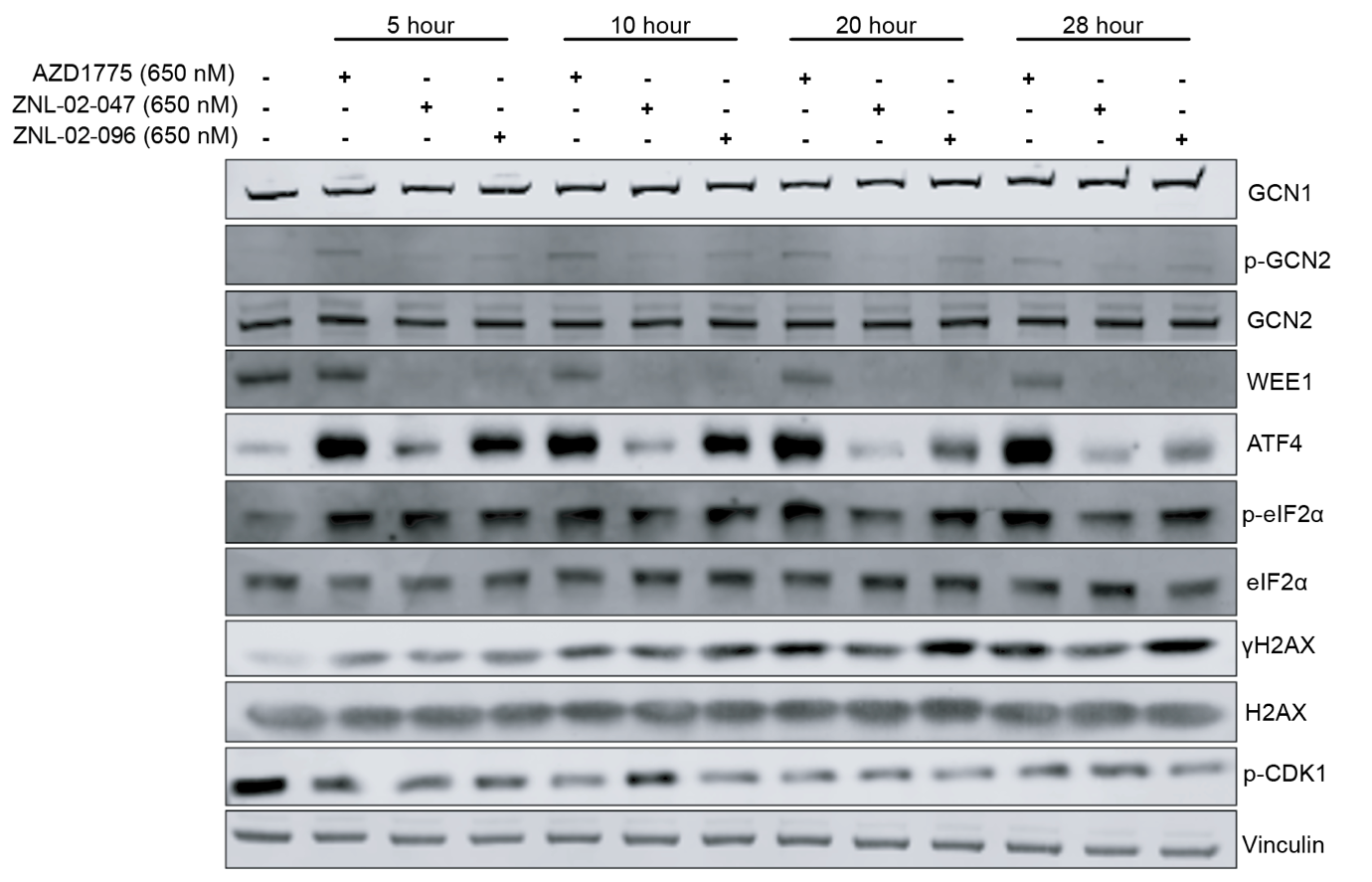
 **b**

**Supplementary Fig.16: A comparison of AZD1775 and PROTAC forms. a** Heatmaps showing resazurin cell viability assays in a 96 well plate format. CC-90009 was treated in combination with AZD1775 or ZNL-02-096 for 72 hours (biological n=4). **b** Western blot showing the comparison of 650 nM AZD1775, 650 nM ZNL-02-047 and 650 nM ZNL-02-096 across multiple timepoints for ISR and γH2AX signals in the RPE *TP53*^-/-^ cell line.

**Supplementary Table 1. List of antibodies used in this study**

| Target | Supplier | Catalogue no. | Application | Dilution | Species |
| --- | --- | --- | --- | --- | --- |
| WEE1 | Cell signaling | CST4936 | Western | 1:500 | Rabbit |
| CDK1 | Abcam | ab32094 | Western | 1:500 | Rabbit |
| CDK1 pY15 | Cell signaling | CST9111 | Western | 1:500 | Rabbit |
| H2AX | Abcam | ab11175 | Western | 1:500 | Rabbit |
| gH2AX pS139 | Merck | 05-636 | Western | 1:1000 | Mouse |
| Vinculin | Abcam | ab219649 | Western | 1:3000 | Rabbit |
| GAPDH | Merck | MAB374 | Western | 1:5000 | Mouse |
| GCN2 | Abcam | ab134053 | Western | 1:500 | Rabbit |
| GCN2 pT899 | Abcam | ab75836 | Western | 1:500 | Rabbit |
| eIF2α | Cell signaling | CST9722 | Western | 1:500 | Rabbit |
| eIF2α pS51 | Cell signaling | CST3597 | Western | 1:500 | Rabbit |
| ATF4 | Cell signaling | CST11815 | Western | 1:500 | Rabbit |
| GSPT1 | Abcam | ab49878 | Western | 1:500 | Rabbit |
| DUT | Proteintech | 11728-1-AP | Western | 1:500 | Rabbit |
| GCN1 | Abcam | ab86139 | Western | 1:500 | Rabbit |
| CDK2 | Cell signaling | CST2546 | Western | 1:500 | Rabbit |
| Puromycin | Merck | MABE343 | Western | 1:2000 | Mouse |
| gH2AX pS139 | Merck | 05-636 | IF | 1:1000 | Mouse |
| ATF4 | Cell signaling | CST11815 | IF | 1:200 | Rabbit |

**Supplementary Table 2. List of sgRNAs and primers used in the study**

| Target | sgRNA sequence (5’-3’) | FW primer (5’-3’) | RV primer (5’-3’) |
| --- | --- | --- | --- |
| sgRNAs used for CRISPRi mediated knockdown | | | |
| GCN2 (#1) | GCAGCGCTGCGCCCAAGGCA |  |  |
| GCN2 (#2) | GGCCCACCGCCGCCCAGGCA |  |  |
| GCN1 (#1) | GGGCGGCGCAGGCAGACCGC |  |  |
| GCN1 (#2) | GGGCGGACACGCAGGTGAGG |  |  |
| GSPT1 (#1) | GAGCTAGCGACAAAGATCCC |  |  |
| GSPT1 (#2) | GCTCGCGACGACGACAGAGG |  |  |
| ALKBH8 (#1) | GGGCGTGCAAGTATCCGCTG |  |  |
| ALKBH8 (#2) | GTGGCCGCGCCCAGGGGAGA |  |  |
| DUT (#1) | GCGAGCGAGGAGACCACCGG |  |  |
| DUT (#2) | GAGGCGAGCGAGGAGACCAC |  |  |
| RRM2 (#1) | GGGACAGGACGGCTGGGACA |  |  |
| FZR1 (#1) | GGCGGTCCCTAATATGGCGG |  |  |
| FZR1 (#2) | GTCCGCGGTCCCTAATATGG |  |  |
| PKMYT1 (#1) | GTCACGGGAGTCCTCCGCCC |  |  |
| PKMYT1 (#2) | GGGGCGTCCGGAACAGTCGA |  |  |
| CDK2 (#1) | GCCGTGGCCCCGGGTCGGGA |  |  |
| CDK2 (#2) | GTGGCGGTCGGGAACTCGGT |  |  |
| CCNE1 (#1) | GGCCGCCAGCGCGGTGTAGG |  |  |
| CCNE2 (#1) | GCCGATCACTTACCACAGGC |  |  |
| CCNA2 (#1) | GACTGAAGTCCGGGAACCCG |  |  |
| AAVS1 (#1) | ACTGTTGACGGCGGCGATGT |  |  |
| AAVS1 (#2) | GCTGATACCGTCGGCGTTGG |  |  |
| Primers used for RT-qPCR | | | |
| GAPDH |  | GTGGTCTCCTCTGACTTCAAC | GGAAATGAGCTTGACAAAGTGG |
| ALKBH8 |  | TGAGGCAAACACCTTGTAACT | CAGCCGTGAGGCTTCTTTAT |
| RRM2 |  | GCAAGCGATGGCATAGTAAATG | AACAGCGGGCTTCTGTAATC |
| FZR1 |  | CTGAGGTTCTGGAACGTCTTTAG | TACCGGATCCTGGTGAAGAG |
| PKMYT1 |  | AGTGGCATGCAACATGGAG | AGCATCATGACAAGGACAGAAC |
| CCNE1 |  | GTGACAGATGGAGCTTGTTC | CATTCAGCCAGGACACAATAG |
| CCNE2 |  | AGTCCAGTGAAGCTGAAGAC | TCCTCCAGCATAGCCAAATAG |
| CCNA2 |  | GATAGGTTCCTGTCTTCCATGT | TACACAAACTCTGCTACTTCTGG |
